# Ecosystem functional response across precipitation extremes in a sagebrush steppe

**DOI:** 10.1101/195594

**Authors:** Andrew T. Tredennick, Andrew R. Kleinhesselink, J. Bret Taylor, Peter B. Adler

## Abstract

**Background:** Precipitation is predicted to become more variable in the western United States, meaning years of above and below average precipitation will become more common. Periods of extreme precipitation are major drivers of interannual variability in ecosystem functioning in water limited communities, but how ecosystems respond to these extremes over the long-term may shift with precipitation means and variances. Long-term changes in ecosystem functional response could reflect compensatory changes in species composition or species reaching physiological thresholds at extreme precipitation levels.

**Methods:** We conducted a five year precipitation manipulation experiment in a sagebrush steppe ecosystem in Idaho, United States. We used drought and irrigation treatments (approximately 50% decrease/increase) to investigate whether ecosystem functional response remains consistent under sustained high or low precipitation. We recorded data on aboveground net primary productivity (ANPP), species abundance, and soil moisture. We fit a generalized linear mixed effects model to determine if the relationship between ANPP and soil moisture differed among treatments. We used nonmetric multidimensional scaling to quantify community composition over the five years.

**Results:** Ecosystem functional response, defined as the relationship between soil moisture and ANPP, was similar among irrigation and control treatments, but the drought treatment had a greater slope than the control treatment. However, all estimates for the effect of soil moisture on ANPP overlapped zero, indicating the relationship is weak and uncertain regardless of treatment. There was also large spatial variation in ANPP within-years, which contributes to the uncertainty of the soil moisture effect. Plant community composition was remarkably stable over the course of the experiment and did not differ among treatments.

**Discussion:** Despite some evidence that ecosystem functional response became more sensitive under sustained drought conditions, the response of ANPP to soil moisture was consistently weak and community composition was stable. The similarity of ecosystem functional responses across treatments was not related to compensatory shifts at the plant community level, but instead may reflect the insensitivity of the dominant species to soil moisture. These species may be successful precisely because they have evolved life history strategies which buffer them against precipitation variability.

## 1 INTRODUCTION

Water availability is a major driver of annual net primary productivity (ANPP) in grassland ecosystems (Huxman et al. 2004; Hsu, Powell, and Adler 2012). Therefore, projected changes in precipitation regimes, associated with global climate change, will impact grassland ecosystem functioning. At any given site, the relationship between ANPP and water availability (e.g., soil moisture) can be characterized by regressing historical observations of ANPP on observations of soil moisture. Fitted functional responses can then be used to infer how ANPP may change under future precipitation regimes (e.g., Hsu, Powell, and Adler 2012). A problem with this approach is that it requires extrapolation if future precipitation falls outside the historical range of variability (Smith 2011; Peters et al. 2012). For example, the soil moisture-ANPP relationship may be linear within the historical range of interannual variation, but could saturate at higher levels of soil moisture. Saturating relationships are actually common (Hsu, Powell, and Adler 2012; Laureano A Gherardi and Sala 2015), perhaps because other resources, like nitrogen, become more limiting in wet years than dry years. Failure to accurately estimate the curvature of the soil moisture-ANPP relationship will lead to over-or underprediction of ANPP under extreme precipitation (Peters et al. 2012).

Another problem with relying on historical ecosystem functional responses to predict impacts of altered precipitation regimes is that these relationships themselves might shift over the long-term. Shifts in species identities and/or abundances can alter an ecosystem’s functional response to water availability because different species have different physiological thresholds. Smith, Knapp, and Collins (2009) introduced the ‘Hierarchical Response Framework’ for understanding the interplay of community composition and ecosystem functioning in response to long-term shifts in resources. In the near term, ecosystem functioning such as ANPP will reflect the physiological responses of individual species to the manipulated resource level. For example, ANPP may initially decline under simulated drought because the community consisted of drought-intolerant species (Hoover, Knapp, and Smith 2014), but functioning may recover over longer time spans as new species colonize or resident species reorder in relative abundance. It is also possible that ecosystem functioning eventually shifts to a new mean state, reflecting the suite of species in the new community (Knapp, Briggs, and Smith 2012).

Experimental manipulations of limiting resources, like precipitation, offer a way to test how ecosystems will respond to resource levels that fall outside the historical range of variability (Avolio et al. 2015; Laureano A. Gherardi and Sala 2015; Knapp et al. 2017). Altering the amount of precipitation over many years should provide insight into the time scales at which water-limited ecosystems respond to chronic resource alteration. We propose four alternative scenarios for the effect of precipitation manipulation on the ecosystem functional response to soil moisture based on the Hierarchical Response Framework (Fig. 1). We define ‘ecosystem functional response’ as the relationship between available soil moisture and ANPP. We focus on soil moisture rather than precipitation because soil moisture is more directly related to plant resource requirements. The four scenarios are based on possible outcomes at the community (e.g., community composition) and ecosystem (e.g., soil moisture-ANPP regression) levels.

**Figure 1.**
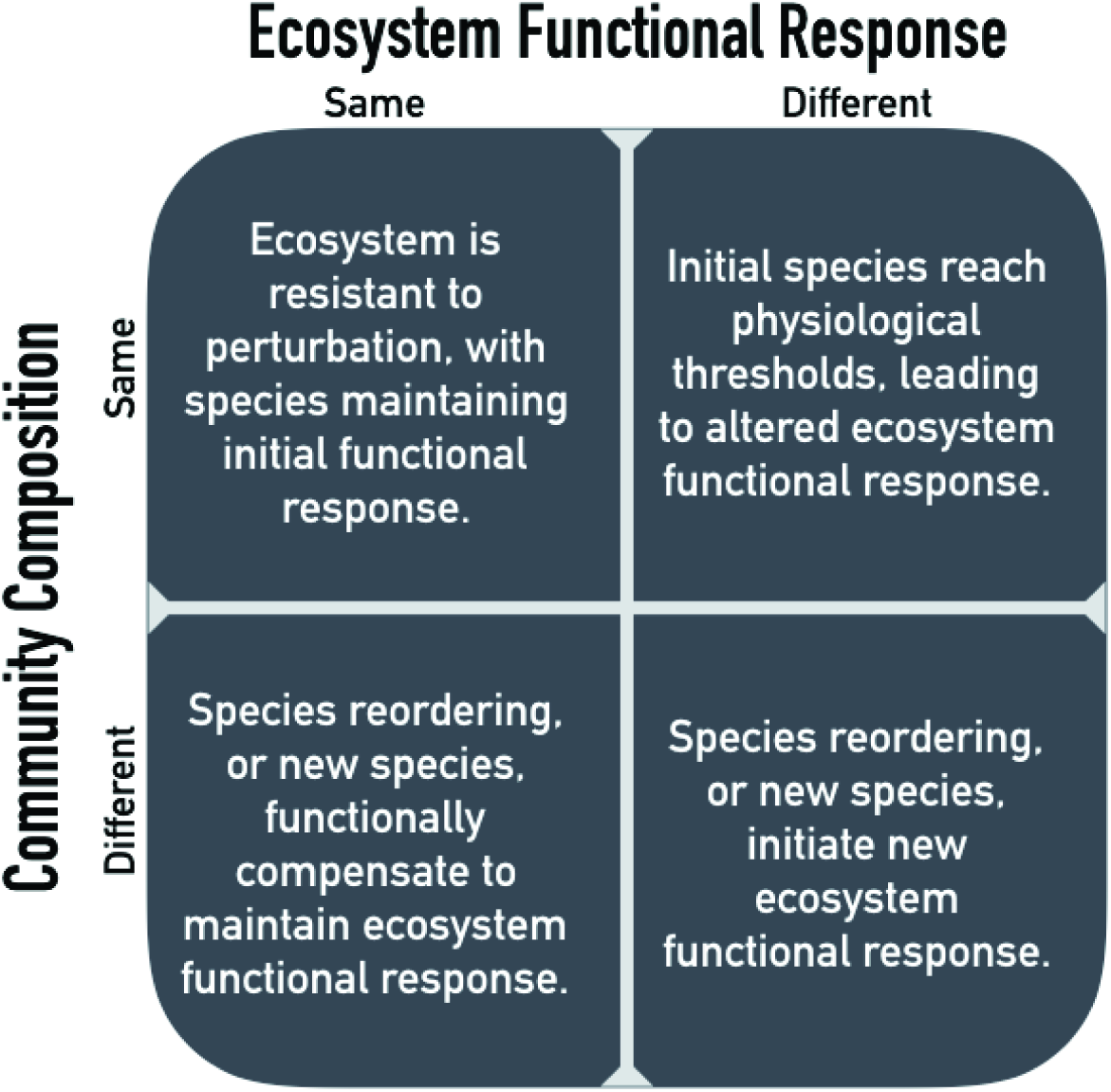
Possible outcomes of chronic resource alteration based on the ‘Hierarchical Response Framework’ (Smith et al. 2009).

First, altered precipitation might have no effect on either ecosystem functional response or community composition (Fig. 1, top left). In this case, changes in ANPP would be well predicted by the current, observed soil moisture-ANPP relationship. This corresponds to the early phases of the Hierarchical Response Framework, where ecosystem response follows the physiological responses of individual species. Second, the ecosystem functional response might change while community composition remains the same (Fig. 1, top right). A saturating soil moisture-ANPP response fits this scenario, where individual species hit physiological thresholds or are limited by some other resource. Third, the ecosystem functional response might be constant but community composition changes (Fig. 1, bottom left). In this case, changes in species’ identities and/or abundances occur in response to altered precipitation levels and species more suited to the new conditions compensate for reduced function of initial residents. Fourth, and last, both ecosystem functional response and community composition could change (Fig. 1, bottom right). New species, or newly abundant species, with different physiological responses completely reshape the ecosystem functional response.

All four outcomes are possible in any given ecosystem, but the time scales at which the different scenarios play out likely differ (Smith, Knapp, and Collins 2009; Wilcox et al. 2016; Knapp et al. 2017). To determine these time scales, we need to amass information on how quickly ecosystem functional responses change in different ecosystems. We also need to understand whether changes at the ecosystem level are driven by community level changes or individual level responses.

To that end, here we report the results of a five-year precipitation manipulation experiment in a sagebrush steppe. We imposed drought and irrigation treatments (approximately ±50%) and measured ecosystem (ANPP) and community (species composition) responses. We focus on how the drought and irrigation treatments affect the relationship between interannual variation in available soil moisture and interannual variation in ANPP, and if community dynamics underlie the ecosystem responses. In particular, we are interested in the consistency of the soil moisture-ANPP relationship among treatments. Is the relationship steeper under the drought treatment at low soil moisture? Does the relationship saturate under the irrigation treatment at high soil moisture? To answer these questions we fit a generalized linear mixed effects model to test whether the regressions differed among treatments. We also analyzed community composition and the sensitivity of ANPP to drought and irrigation treatments over time, allowing us to place our experimental results within the framework of our scenarios (Fig. 1).

## 2 METHODS

### 2.1 Study Area

We conducted our precipitation manipulation experiment in a sagebrush steppe community at the USDA-ARS Sheep Experiment Station (USSES) near Dubois, Idaho (44.2° N, 112.1° W), 1500 m above sea level. The plant community is dominated by the shrub *Artemisia tripartita*, the perennial forb *Balsamorhiza sagittata*, and three perennial bunchgrasses, *Pseudoroegneria spicata, Poa secunda*, and *Hesperostipa comata* (see Appendix 1 for rank abundance curves). During the period of our experiment (2011 - 2016), mean annual precipitation was 250 mm year^−1^ and mean monthly temperature ranged from −5.2°C in January to 21.8°C in July. Between 1926 and 1932, range scientists at the USSES established 26 permanent 1 m^2^ quadrats to track vegetation change over time. In 2007, we located 14 of the original quadrats in permenanent livestock exclosures, which we used as control plots (i.e. ambient precipitation) in the experiment described below. We used the original plots as our controls because collecting demographic data is time consuming and it was already being collected in these plots for other studies.

In spring 2011, we established 16 new 1 m^2^ plots located in the largest exclosure at USSES, which also contained six of our control plots. We avoided areas on steep hill slopes, areas with greater than 20% cover of bare rock, and areas with greater than 10% cover of the shrubs *Purshia tridentata* and/or *Amelanchier utahensis.* We established the new plots in pairs and randomly assigned each plot in a pair to receive the “drought” or “irrigation” treatment. Thus, our experiment consisted of *n* = 14 control plots, *n* = 8 irrigation plots, and *n* = 8 irrigation plots, for a total of 30 plots.

### 2.2 Precipitation Experiment

Drought and irrigation treatments were designed to decrease and increase the amount of ambient precipitation by 50%. To achieve this, we used a system of rain-out shelters and automatic irrigation (Gherardi and Sala 2013). The rain-out shelters consisted of transparent acrylic shingles 1-1.5 m above the ground that covered an area of 2.5 × 2 m. The shingles intercepted approximately 50% of incoming rainfall, which was channeled into 75 liter containers. Captured rainfall was then pumped out of the containers and sprayed on to the adjacent irrigation plot via two suspended sprinklers. Pumping was triggered by float switches once water levels reached about 20 liters. We disconnected the irrigation pumps each October and reconnected them each April. The rain-out shelters remained in place throughout the year.

We monitored soil moisture in four of the drought-irrigation pairs using Decagon Devices (Pullman, Washington) 5TM and EC-5 soil moisture sensors. We installed four sensors around the edges of each 1x1 m census plot, two at 5 cm soil depth and two at 25 cm soil depth. We also installed four sensors in areas nearby the four selected plot pairs to measure ambient soil moisture at the same depths. Soil moisture measurements were automatically logged every four hours. We coupled this temporally intensive soil moisture sampling with spatially extensive readings taken with a handheld EC-5 sensor at six points within all 16 plots and associated ambient measurement areas. These snapshot data were collected on 06-06-2012, 04-29-2015, 05-07-2015, 06-09-2015, and 05-10-2016^1^.

Analyzing the response to experimental treatments was complicated by the fact that we did not directly monitor soil moisture in each plot on each day of the experiment. Only a subset of plots were equipped with soil moisture sensors, and within those plots, one or more of the sensors frequently failed to collect data. Therefore, we used a statistical model to estimate average daily soil moisture values for the ambient, drought, and irrigation treatments during the course of the experiment.

We first averaged the observed soil moisture readings for each day (*d*)and plot (*i*), *x*_*i,d*_. Experimental plots were located in pairs, with each group (*g*)containing a drought and irrigated plot. In addition, a nearby area outside the drought or irrigated plots was monitored for local ambient soil moisture conditions. Within each group of plots we standardized the irrigation and drought effects on soil moisture relative to the ambient soil moisture conditions. Specifically, we subtracted the ambient daily soil moisture from the soil moisture measured within the drought and irrigation plots within each group and then divided by the standard deviation of daily soil moisture values measured in the ambient conditions (Δ*x*_*g,d*,trt_. = (*x*_*g,d*,trt_. − *x*_*g,d*,ambient_)/s.d.(*x*_*g,d*,ambient_)) where Dx is the standardized treatment effect and *x*_*g,d*,trt_ is the raw soil moisture measure for plot group *g* on day *d* and treatment *trt*. These transformations ensured that the treatment effects in each plot were appropriately scaled by the local ambient conditions within each plot group.

We then modeled these daily deviations (Δ*x*_*g,d*,trt._) from ambient conditions using a linear mixed effects model with independent variables for treatment (irrigation or drought), season (winter, spring, summer, fall), rainfall, and all two-way interactions. Using the local daily weather station data, we recorded rainy days as any day with measureable precipitation or the day after such a day and with average temperatures above 3°C (to exclude days with snowfall). We fit the model using the `lme4∷lmer()` function (Bates et al. 2015) in R (R Core Team 2016), with random effects for plot group and date. We weighted observations by the number of unique sensors or spot measurements that were taken in each plot on that day. We then used the model to predict the deviations from ambient conditions produced in the treated plots on each day of the experiment. We added these predicted deviations to the average daily ambient soil moisture to generate predictions for daily soil moisture in all of the treated plots: 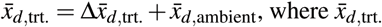 is the average predicted daily soil moisture in the treated plots and 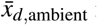 ambient is the daily ambient soil moisture averaged across all control plots. We could only predict soil moisture in the treated plots on days for which we took at least one ambient soil moisture measurement. This procedure allowed us to predict daily soil moisture conditions for all plots, even on those days when some of our direct treatment measurements were missing due to malfunction of the sensors or data loggers. See Kleinhesselink (2017) for more details on our approach to estimating soil moisture.

Following the above procedure, we still lacked soil moisture data for March 2012 (observations did not start until April 2012) and for a string of days in 2013 during which the soil moisture probes failed to take readings. To fill in these gaps, we used a version of the SOILWAT soil moisture model (Sala, Lauenroth, and Parton 1992) that has been specifically designed for semiarid shrublands and grasslands (Bradford, Schlaepfer, and Lauenroth 2014). The model was parameterized using generic sagebrush steppe parameters and local soil texture, soil bulk density, and precipitation data (Kleinhesselink 2017). We used the locally parameterized SOILWAT model to generate daily soil moisture predictions for the duration of the experiment, but only used SOILWAT predictions where there were gaps in our data (Fig. 2B and Appendix 2). SOILWAT predicted daily soil moisture under ambient conditions similar to our control plots. We applied the same statistical model and procedure described above to estimate soil moisture in drought and irrigation plots based on control plot conditions.

**Figure 2.**
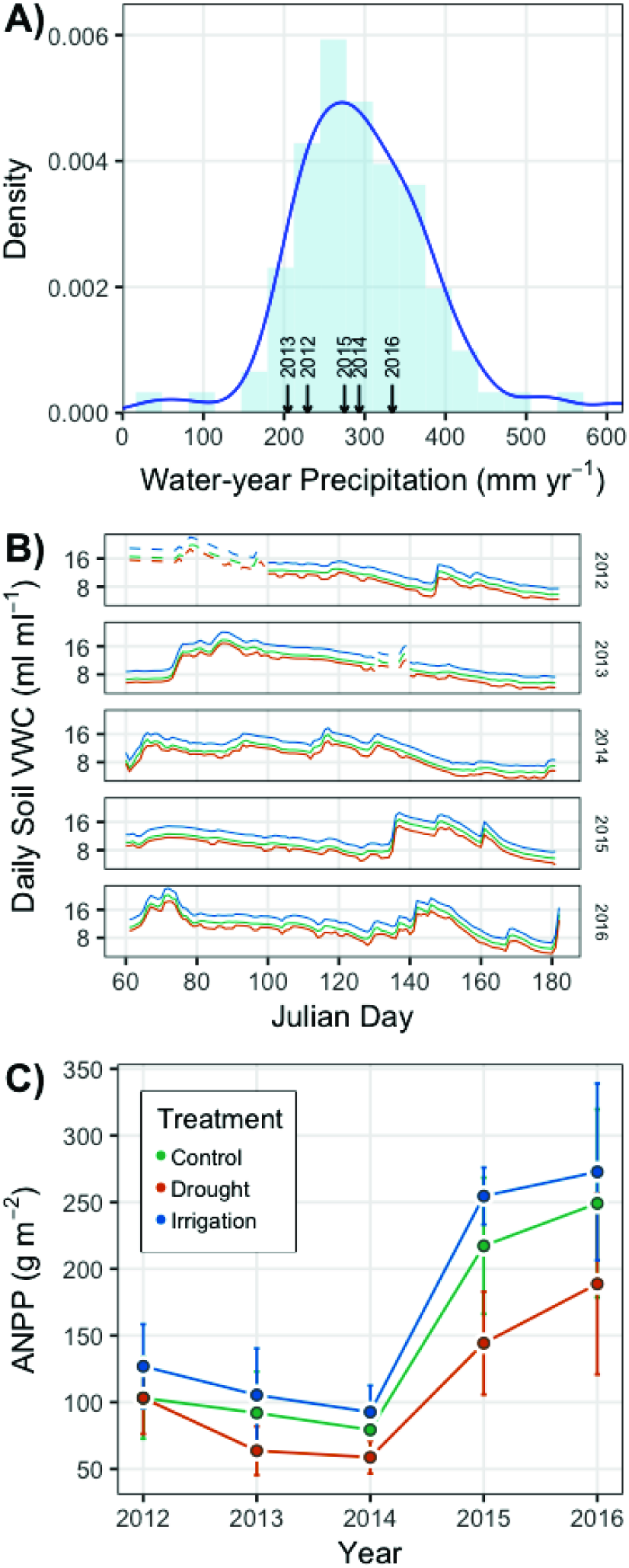
(A) Probability density of historical water-year (Oct.-Sept.) precipitation from 1927-2016, with the years of the experiment shown by arrows on the *x*-axis. (B) Statistically estimated (solid lines) and SOILWAT-generated (dashed lines) daily soil volumetric water content (VWC) in each of the three treatments during the course of the experiment. We used the estimates from our statistical model, with gaps filled in by SOILWAT predictions, to calculate cumulative March-June VWC as used in our analysis of treatment effects on ecosystem functional response. (C) Mean (points) ANPP and its standard deviation (error bars) for each year of the experiment. Colors in panels B and C identify the treatment, as specified in the legend of panel C.

### 2.3 Data Collection

We estimated aboveground net primary productivity (ANPP) using a radiometer to relate ground reflectance to plant biomass (Byrne et al. 2011). We recorded ground reflectance at four wavelengths, two associated with red reflectance (626 nm and 652 nm) and two associated with near-infrared reflectance (875 nm and 859 nm). At each plot in each year, we took four readings of ground reflectances at the above wavelengths. We also took readings in 12 (2015), 15 (2012, 2013, 2014), or 16 (2016) calibration plots adjacent to the experimental site, in which we harvested all aboveground biomass produced in the current year (we excluded litter and standing dead material), dried it to a constant weight at 60°C, and weighed it to estimate ANPP. We made radiometer measurements and harvested at peak green biomass each year, typically in late June.

For each plot and year, we averaged the four readings for each wavelength and then calculated a greenness index based on the same bands used to calculate NDVI using the MODIS (Moderate Resolution Imaging Spectroradiometer) and AVHRR (Advanced Very High Resolution Radiometer) bands for NDVI. We regressed the greenness index against the dry biomass weight from the calibration plots to convert the greenness index to ANPP. We fit regressions to a MODIS-based index and an AVHRR-based index for each year and retained the regression with the better fit based on *R*^2^ values. We then predicted ANPP using the best regression equation for each year (Appendix 3). Our results do not change when we analyze the soil moisture-NDVI relationship instead of the soil moisture-ANPP relationship (Appendix 5).

Species composition data came from two sources: yearly census maps for each plot made using a pantograph (Hill 1920) and yearly counts of annual species in each plot. From these sources, we determined the density of all annuals and perennials forbs, the basal cover of perennial grasses, and the canopy cover of shrubs. We made a large plot-treatment-year by species matrix, where columns were filled with either cover or density, depending on the measurement made for the particular species. We standardized the values in each column so we could directly compare species quantified with different metrics (density, basal cover, and canopy cover). This puts all abundance values on the same scale, meaning that common and rare species are weighted equally. Assuming that rare species will respond to treatments more than common ones, our approach is biased towards detecting compositional changes. We limited our analysis of community data to observations from the permanent exclosure containing our drought and irrigation treatments (see **Community composition over time**). This exclosure included six of the control plots and all of the treatment plots, for a total of 22 plots. The other eight control plots are in other pastures. Including these plots would add spatial variation in composition, complicating our goal of describing temporal trends in composition.

### 2.4 Data Analysis

#### 2.4.1 Ecosystem functional response

Our main goal was to test whether the relationship between ANPP and soil moisture differed among the drought, control, and irrigation treatments. Based on our own observations and previous work at our study site (Blaisdell 1958; Dalgleish et al. 2011; Adler, Dalgleish, and Ellner 2012), we chose to use cumulative volumetric water content from March through June as our metric of soil moisture (hereafter referred to as ‘VWC’). We fit a generalized linear mixed effects regression model with log(ANPP) as the response variable and VWC and treatment as fixed effects. Plot and year of treatment were included as random effects to account for non-independence of observations, as described below. We log-transformed ANPP to reduce heteroscedasticity. Both log(ANPP) and VWC were standardized to have mean 0 and unit variance before fitting the model 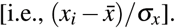.

Our model is defined as follows:

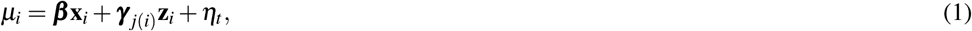

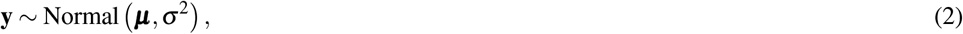

where μ_*i*_ is the deterministic prediction from the regression model for observation *i*, which is associated with plot *j* and treatment year *t*. **β** is the vector of coefficients for the fixed effects in the design matrix **X**. Each row of the design matrix represents a single observation (**x**_*i*_) and is a vector with the following elements: 1 for the intercept, a binary 0 or 1 if the treatment is “drought”, a binary 0 or 1 if the treatment is “irrigation”, the scaled value of VWC, binary “drought” value times VWC, and binary “irrigation” value times VWC. Thus, our model treats “control” observations as the main treatment and then estimates intercept and slope offsets for the “drought” and “irrigation” treatments. We use our model to test two statistical hypotheses based on the questions in our Introduction:

**H1**. The coefficient for drought?VWC is positive and different from zero.

**H2**. The coefficient for irrigation?VWC is negative and different from zero.

These hypotheses are based on evidence that precipitation-ANPP relationships often saturate with increasing precipitation (Hsu, Powell, and Adler 2012; Laureano A Gherardi and Sala 2015).

We include two random effects to account for the fact that observations within plots and years are not independent. Specifically, we include plot-specific offsets (***γ***) for the intercept and slope terms and year-specific intercept offsets (*η*_*t*_). The covariate vector **z**_*i*_ for each observation i has two elements: a 1 for the intercept and the scaled value of VWC for that plot and year. The plot-specific coefficients are modeled hierarchically, where plot level coefficients are drawn from a multivariate normal distribution with mean 0 and a variance-covariance structure that allows the intercept and slope terms to be correlated:

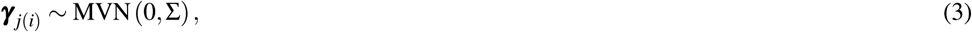

where Σ is the variance-covariance matrix and *j(i*) reads as “plot *j* associated with observation *i*”. The random year effects (**η**) are drawn from a normal prior with mean 0 and standard deviation s_year_, which was drawn from a half-Cauchy distribution. We used a vague prior distribution for each *β*: ***β*** ~ Normal(0,5). A full description of our model and associated R (R Core Team 2016) code is in Appendix 4.

We fit the model using a Bayesian approach and obtained posterior estimates of all unknowns via the No-U-Turn Hamiltonian Monte Carlo sampler in Stan (Stan Development Team 2016b). We used the R package ‘rstan’ (Stan Development Team 2016a) to link R (R Core Team 2016) to Stan. We obtained samples from the posterior distribution for all model parameters from four parallel MCMC chains run for 10,000 iterations, saving every 10^th^ sample. Trace plots of all parameters were visually inspected to ensure well-mixed chains and convergence. We also made sure all scale reduction factors (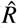 values) were less than 1.1 (Gelman and Hill 2009).

#### 2.4.2 Sensitivity of ANPP over time

Our data do not allow us to directly test whether the ecosystem functional response to precipitation may have changed over time since the start of the experiment in each of the treatments. This is because we lack sufficient within-year and within-treatment variation of soil moisture (i.e., within a year and treatment each plot shares the same value of VWC). However, we did use a separate analysis to test whether the the sensitivity of ANPP to treatment changed over time.

We define ‘sensitivity’ as 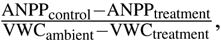, following Wilcox et al. (2017). This metric of sensitivity scales the difference between ANPP in the treated and control plots by the change in soil moisture in the treated plots. Because our treated plots are not directly paired with individual control plots, we compare ANPP in each treated plot to the average ANPP of the control plots in each year. After calculating sensitivity for each drought and irrigation plot in each year, we regressed sensitivity against year of treatment using the `lm()` function in R (R Core Team 2016). This analysis also allows us to link particularly sensitive treatment-years to changes in community composition.

#### 2.4.3 Community composition over time

We used nonmetric multidimensional scaling (NMDS) based on Bray-Curtis distances to identify temporal changes in community composition among treatments. We first calculated Bray-Curtis distances among all plots for each year of the experiment and then extracted those distances for use in the NMDS. Some values of standardized species’ abundances were negative, which is not allowed for calculating Bray-Curtis distances. We simply added ‘2’ to each abundance value to ensure all values were greater than zero. We plotted the first two axes of NMDS scores to see if community composition overlapped, or not, among treatments in each year. We used the `vegan∷metaMDS()` function (Oksanen 2016) to calculate Bray-Curtis distances and then to run the NMDS analysis. We used the `vegan∷adonis()` function (Oksanen 2016) to perform permutational multivariate analysis of variance to test whether treatment plots formed distinct groupings. To test whether treatment plots were equally dispersed, or not, we used the `vegan∷betadisper()` function (Oksanen 2016).

We conducted the above analysis for all species, and then conducted a separate analysis for annual species only (Appendix 1). Annual species have shorter life spans, so conducting a separate analysis allowed us to test whether we might find stronger evidence for community responses to altered precipitation when we focus on short-lived species. Given the dominance of perennial species in our system (Appendix 1), shifts in the annual plant community could be difficult to detect in the analysis of the full community.

#### 2.4.4 Reproducibility

All R code and data necessary to reproduce our analysis has been archived on Figshare (*link here after acceptance*) and released on GitHub (https://github.com/atredennick/usses_water/releases/v0.1). We also include annotated Stan code in our model description in Appendix 4.

## 3 RESULTS

Ambient precipitation and soil moisture were variable over the five years of the study (Fig. 2A-B). ANPP varied from a minimum of 77.4 g m^−2^ in 2014 to a maximum of 239.3 g m^−2^ in 2016 when averaged across treatments (Fig. 2C). ANPP was slightly higher in irrigation plots (on average 15% higher) and slightly lower in drought plots (on average 25% lower) relative to control plots (Fig. 2C), corresponding to observed and estimated soil volumetric water content (VWC) differences among treatments (Fig. 2B). March-June VWC in drought plots was 12% less than in control plots on average, and March-June VWC in irrigation plots was 19% higher than in control plots on average across the years of the experiment. The differences in soil VWC indicate our treatment infrastructure was successful. ANPP was highly variable across plots within years, as indicated by the large and overlapping standard deviations (Fig. 2C).

Cumulative March-June soil moisture had a weak positive effect on ANPP (Table 1; Fig. 3B). The effect of soil moisture for each treatment is associated with high uncertainty, however, with 95% Bayesian credible intervals that broadly overlap zero (Table 1). Although the parameter estimates for the effect of soil moisture overlap zero, the posterior distributions of the slopes all shrank and shifted to more positive values relative to the prior distributions (Fig. A3-2), which indicates the data did influence parameter estimates. Ecosystem functional response was similar among treatments (Table 1; Fig. 3B), but there is evidence that the slope for the drought treatment is greater than the slope for the control treatment. This evidence comes from interpreting the posterior distributions of the slope offsets for the treatments. From these distributions, we calculate a 42% one-tailed probability that the estimate is less than zero for the irrigation treatment and a 100% one-tailed probability that the estimate is greater than zero for the drought treatment (Fig. 3A, right panel).

**Table 1.**
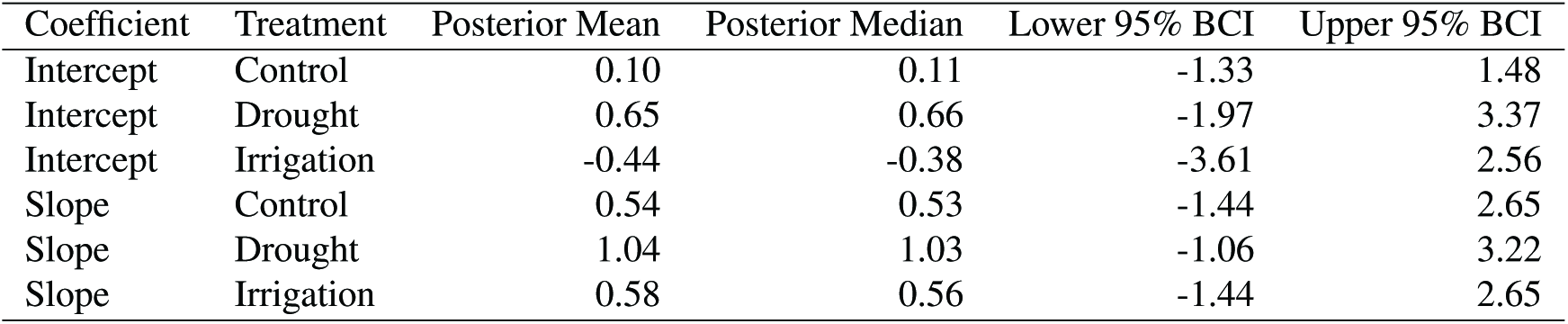
Summary statistics from the posterior distributions of coefficients for each treatment (*β* coefficients in equation 1). The ‘Intercept’ and ‘Slope’ summaries reported here for drought and irrigation are from the posterior distributions of the intercept and slope for the control treatment plus the offsets for each treatment. Posterior distributions of the offsets are in Figure 3A.

**Figure 3.**
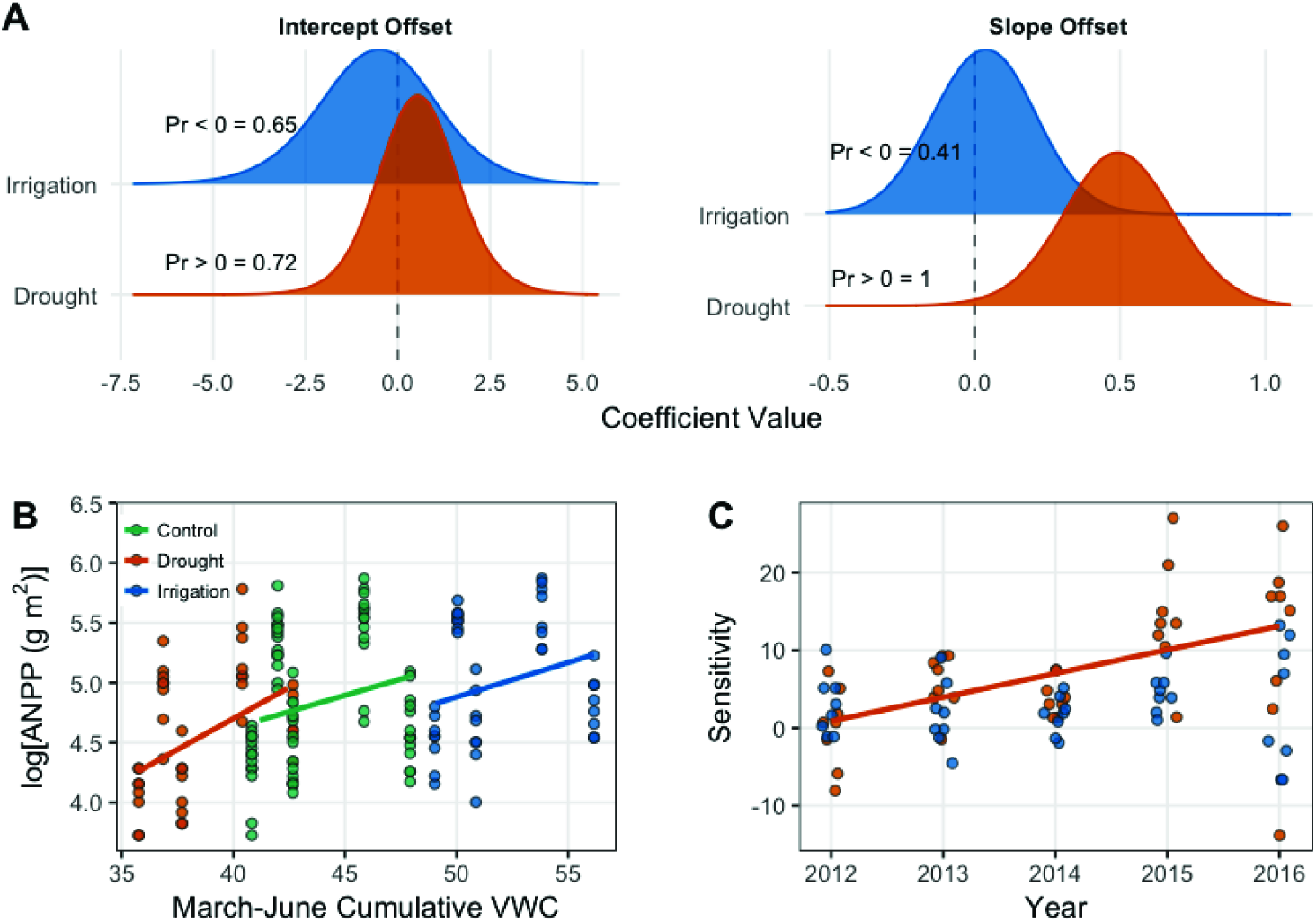
Results from the generalized linear mixed effects model (A-B) and the sensitivity analysis (C). (A) Posterior distributions for the effects of drought and irrigation on the intercept and slope of the productivity-soil moisture relationship. Treatment effects show the difference between the coefficients estimated in the treated plots and the control plots. Probabilities (“Pr < / = 0 =”) for each coefficient indicate the one-tailed probability that the coefficient is less than or greater than zero, depending on the median of the intercept offset distributions or our specific hypothesis for the slope offsets. The posterior densities were smoothed for visual clarity by increasing kernel bandwidth by a factor of five. (B) Scatterplot of the data and model estimates shown as solid lines. Model estimates come from treatment level coefficients (colored lines). Note that we show log(ANPP) on the y-axis of panel B; this same plot can be seen on the arithmetic scale in supporting material Fig. A2-1. (C) Regression of sensitivity against time for each treatment. Each point represents the sensitivity of ANPP in a plot relative to the mean of the control plots in that year. Only the significant regression for the drought treatment is plotted (P = 0.0007).

Sensitivity of ANPP to irrigation was constant over the course of our experiment (Fig. 3C). Sensivitity of ANPP to drought, however, grew over time (P = 0.0007; Fig. 3C).

Community composition was similar among treatments. The multidimensional space of community composition overlapped among treatments in all years and was equally dispersed in all years (Table 2; Fig. 4). Community composition was also remarkably stable over time, with no evidence of divergence among treatments (Table 2; Fig. 4). There were also no changes in the dominant species over time in any treatment (Appendix 1). Analyzing annual species on their own produced similar results. There is some evidence that the annual community changed in response to treatment in two years (Table A1-1; Fig. A1-5), but these responses appear to come from very rare species since our analysis weights common and rare species equally (Fig. A1-4). By definition, these rare species contribute little to ANPP and thus any compositional changes have little influence on ecosystem functional response.

**Table 2.** Results from statistical tests for clustering and dispersion of community composition among precipitation treatments. ‘adonis’ tests whether treatments form unique clusters in multidimensional space; ‘betadisper’ tests whether treatments have similar dispersion. For both tests, P values greater than 0.05 indicate there is no support for the hypothesis that the treatments differ.

| Year | Test | n | d.f. | $F$ | $P$ |
| --- | --- | --- | --- | --- | --- |
| 2011 | adonis | 22 | 2 | 1.04 | 0.40 |
| 2011 | betadisper | 22 | 2 | 2.22 | 0.14 |
| 2012 | adonis | 22 | 2 | 1.09 | 0.32 |
| 2012 | betadisper | 22 | 2 | 0.21 | 0.81 |
| 2013 | adonis | 22 | 2 | 1.26 | 0.09 |
| 2013 | betadisper | 22 | 2 | 0.43 | 0.66 |
| 2014 | adonis | 22 | 2 | 0.96 | 0.54 |
| 2014 | betadisper | 22 | 2 | 0.36 | 0.70 |
| 2015 | adonis | 22 | 2 | 1.08 | 0.30 |
| 2015 | betadisper | 22 | 2 | 1.96 | 0.17 |
| 2016 | adonis | 22 | 2 | 1.12 | 0.27 |
| 2016 | betadisper | 22 | 2 | 0.27 | 0.76 |

**Figure 4.**
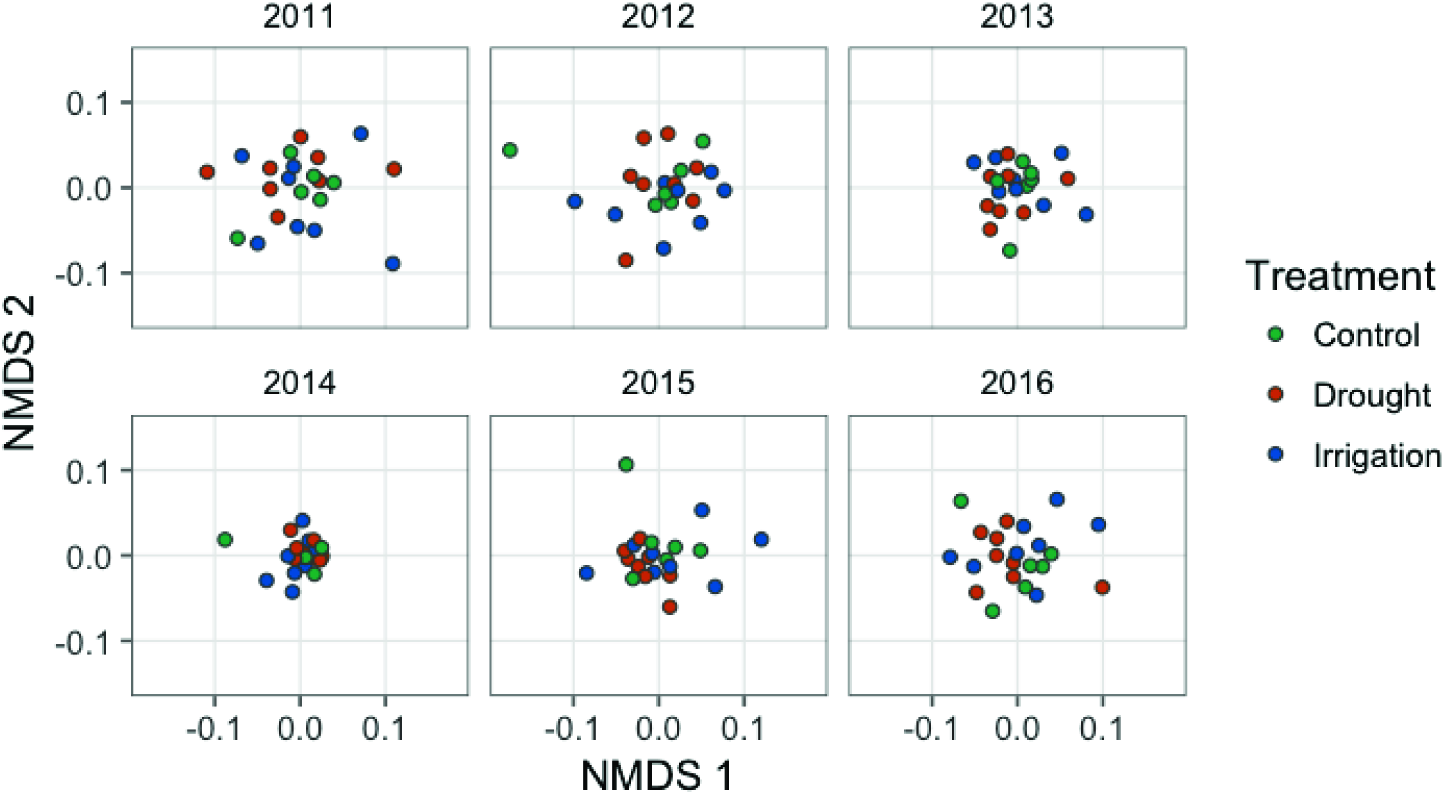
Nonmetric multidimensional scaling scores representing plant communities in each plot, colored by treatment. Each point represents a plot and the axes represent species composition.

## 4 DISCUSSION

Ecosystem response to a new precipitation regime depends on the physiological responses of constituent species and the rate at which community composition shifts to favor species better able to take advantage of, or cope with, new resource levels (Smith, Knapp, and Collins 2009). Previous work has shown that community compositional shifts can be either rapid, on the order of years (Hoover, Knapp, and Smith 2014), or slow, on the order of decades (Knapp, Briggs, and Smith 2012; Wilcox et al. 2016). A lingering question is how the time scales of ecosystem response and community change vary among ecosystems. Precipitation manipulation experiments can help answer this question, especially if they push water availability outside the historical range of variability for long periods.

We found that ecosystem functional response under chronic drought was different from the control treatment, but community composition remained unchanged (Fig. 3A, Fig. 4), representing the top right scenario in Fig. 1. The increase in the slope of the VWC × productivity relationship in the drought plots indicates increased sensitivity to water availability under chronic drought. A strict interpretation of this result implies that if average soil moisture were pushed consistently lower than currently observed ambient conditions, there would be a stronger relationship between precipitation and productivity in this system.

However, we do not want to over-interpret the significance of the slope offsets given that the overall slopes of the VWC × productivity relationship, not just the offsets, were similar among treatments (Table 1). We therefore conclude that ecosystem functional response is consistent (similar values) and weak (all broadly overlapping zero) across all precipitation treatments.

The similarity of ecosystem functional response (Fig. 3) and community composition (Fig. 4) among treatments is surprising because grasslands generally, and sagebrush steppe specifically, are considered water-limited systems. For example, Huxman et al. (2004) and Knapp et al. (2015) showed that semi-arid sites are more sensitive to drought than mesic sites, and Wilcox et al. (2017) found that semi-arid sites are particularly sensitive to irrigation treatments. Based on these findings, we expected ecosystem functional response, community composition, or both to change under our treatments. Why did our treatments fail to induce ecosystem or community responses? We can think of three reasons. Two are limitations of our study, and one invokes the life history traits of the species at our study site.

First, our precipitation manipulations may not have been large enough to induce a response. A 50% decrease in any given year may not be abnormal given our site’s historical range of variability (Knapp et al. 2017). We cannot definitively rule out this possibility, but we have reason to believe our treatments should have been large enough. Using the methods described by Lemoine et al. (2016), we calculated the percent reduction and increase of water-year precipitation necessary to reach the 1% and 99% extremes of the historical precipitation regime at our site (Fig. A5-1). The 1% quantile of water-year precipitation at our site is 78 mm, a 26% reduction from the mean, and the 99% quantile is 545 mm, a 84% increase from mean growing season precipitation (Appendix 6). Thus, our drought treatment represented extreme precipitation amounts, and that is the treatment where we observed a small effect on the slope between soil moisture and ANPP (Fig. 3A). The irrigation treatment may not have been extreme enough to induce a response.

Second, ANPP at our site is likely influenced by additional factors, not only the cumulative March-June soil moisture covariate we included in our statistical model. For example, La Pierre et al. (2016) found that site-scale ANPP is better predicted by nutrient availability than precipitation. Moreover, temperature can impact ANPP directly (Epstein, Lauenroth, and Burke 1997) and by exacerbating the effects of soil moisture (De Boeck et al. 2011). Measurements of soil moisture likely contain a signal of temperature, through its impact on evaporation and infiltration, but the measurements will not capture the direct effect of temperature on metabolic and physiological processes. We also did not redistribute snow across our treatments in the winter, and snow melt may spur early spring growth. Failure to account for potentially important covariates could explain the within-year spread of ANPP (Fig 2C, Fig. 3B) and the resulting uncertain relationship we observed between soil moisture and ANPP across all treatments (Table 1, Fig. A3-2).

Third, the life history traits of the dominant species, which largely determine ecosystem function (Smith and Knapp 2003), in our study ecosystem may explain the weak and uncertain effect of soil moisture on ANPP (Table 1, Fig. 3). Species that live in variable environments, such as cold deserts, must have strategies to ensure long-term success as conditions vary. One strategy is bet hedging, where species forego short-term gains to reduce the variance of long-term success (Seger 1987). In other words, species follow the same conservative strategy every year, designed to minimize losses during unfavorable periods. The dry and variable environment of the sagebrush steppe has likely selected for bet hedging species that can maintain function at low water availability and have weak responses to high water availability. In so doing, the dominant species in our ecosystem avoid “boom and bust” cycles, which may explain the weak effect of soil moisture on ANPP (i.e., the Bayesian credible intervals for the slopes overlapping zero; Table 1).

Another strategy to deal with variable environmental conditions is avoidance, which would also result in a consistent ecosystem functional response between drought and control treatments. For example, many of the perennial grasses in our focal ecosystem avoid drought stress by growing early in the growing season (Blaisdell 1958, A.R. Kleinhesselink, personal observation). Furthermore, the dominant shrub in our focal ecosystem, *Artemisia tripartita*, has access to water deep in the soil profile thanks to a deep root system (Kulmatiski et al. 2017). The dominance of our site by the shrub *A. tripartita* (Appendix 1) may explain why our results do not conform to broader patterns of grassland sensitivity to precipitation manipulations (e.g., Huxman et al. 2004; Knapp et al. 2015; Wilcox et al. 2017).

We found that ANPP became more sensitive to the drought treatment over time (Fig. 3C). We interepret this increase in sensitivity as the effect of cumulative impacts of the drought on dominant species. Plants may not have shown a large response in terms of ANPP in the first years of the experiment if they could grow from stored carbohydrate reserves or if they abstained from flowering and reproduction. As the drought progressed, these same plants may have have started to shrink or die. Given the long-lived perennial species at this site, this increase in sensitivity may indicate a larger future change in community composition and ecosystem functional response.

## 5 CONCLUSIONS

Our results suggest that the species in our focal plant community are insensitive to to changes in precipitation regime, at least over the five years of our experiment. Such insensitivity can buffer species against precipitation variability in this semi-arid ecosystem, making them successful in the long run. Longer, chronic precipitation alteration might reveal plant community shifts that we did not observe (e.g., Wilcox et al. 2016), and the increased sensitivity of ANPP to drought in the final years of our experiment may portend such a shift. Likewise, a long-term increase in water availability could allow species that do not bet hedge to gain prominence and dominate the ecosystem functional response. The length of the perturbation may be especially relevant in our focal ecosystem where the perennial species are long-lived, meaning compositional turnover may take many more years than we report on here.

## 6 ACKNOWLEDGEMENTS

We thank the many summer research technicians who collected the data reported in this paper and the USDA-ARS Sheep Experiment Station (Dubois, ID) for facilitating work on their property. We also thank Susan Durham for clarifying our thinking on the statistical analyses and Kevin Wilcox for helpful discussions on analyzing community composition data. John Bradford and Caitlin Andrews generously provided SOILWAT model output. Two anonymous reviewers and Elsie Denton provided thoughtful reviews that improved our paper.

## 8 AUTHOR CONTRIBUTIONS

1. Andrew T. Tredennick collected data, analyzed the data, wrote the paper, prepared figures and/or tables, reviewed drafts of the paper.
2. Andrew R. Kleinhesselink conceived and designed the experiments, performed the experiments, collected data, analyzed the data, reviewed drafts of the paper.
3. J. Bret Taylor contributed reagents/materials/analysis tools, reviewed drafts of the paper.
4. Peter B. Adler conceived and designed the experiments, performed the experiments, collected data, analyzed the data, reviewed drafts of the paper.

## 9 SUPPLEMENTAL INFORMATION

**Appendix 1**. Additional information on plant community composition and dynamics, Table A1-1, Fig. A1-1, Fig. A1-2, Fig. A1-3, Fig. A1-4, and Fig. A1-5.

**Appendix 2**. Additional details on soil moisture modeling with SOILWAT and Fig. A2-1.

**Appendix 3**. Additional methods and information on estimating aboveground net primary productivity, Table A3-1, and Table A3-2.

**Appendix 4**. Details of the hierarchical Bayesian model, Fig. A4-1, Fig. A4-2, and Fig. A4-3.

**Appendix 5**. Results from analysis of NDVI without conversion to ANPP, Fig. A5-1.

**Appendix 6**. Details on analysis of precipitation historical range of variability and Fig. A6-1.

## Appendix 1 A.T. Tredennick, A.R. Kleinhesselink, J.B. Taylor & P.B. Adler “Consistent ecosystem functional response across precipitation extremes in a sagebrush steppe PeerJ

**Section A1.1 Details on plant community structure**

Here we provide more details on the plant community in terms of dominance and rarity. Averaging across time, *Artemisia tripartita* and *Balsamorhiza sagittata* are the two most dominant species in each treatment. Combined, these two species represent 28% of total cover in control plots, 25% of total cover in drought plots, and 25% of total cover in irrigation plots. Four to five species dominate the community in general (Figure A1-1), indicating a high level of dominance in this plant community.

We also conducted our community composition analysis with only annual species. Annual species are shorter-lived than the perennial species in our community, so they may respond more quickly to alterations of precipitation. In general, our results for annual species conform to the results from the full community analysis in the main text. Annual plant community composition is relatively stable through time (Fig. A1-5) and in most years there is no evidence that treatment differentiates community composition (Table A1-1). Note that in some years the `vegan∷metaMDS()` returned unreliable estimates of Bray-Curtis distances for the annual community because of lack of sufficient data (i.e., many annual species with 0 abundance).

**Section A1.2 Tables**

**Table A1-1.** Results from statistical tests for clustering and dispersion of community composition among precipitation treatments for annual species only. ‘adonis’ tests whether treatments form unique clusters in multidimensial space; ‘betadisper’ tests whether treatments have similar dispersion. For both tests, *P* values greater than 0.05 indicate there is no support that the treatments differ.

| Year | Test | n | d.f. | $F$ | $P$ |
| --- | --- | --- | --- | --- | --- |
| 2011 | adonis | 22 | 2 | 1.09 | 0.38 |
| 2011 | betadisper | 22 | 2 | 5.61 | 0.01 |
| 2012 | adonis | 19 | 2 | 2.67 | 0.02 |
| 2012 | betadisper | 19 | 2 | 0.95 | 0.41 |
| 2013 | adonis | 22 | 2 | 1.71 | 0.10 |
| 2013 | betadisper | 22 | 2 | 1.03 | 0.38 |
| 2014 | adonis | 22 | 2 | 1.37 | 0.15 |
| 2014 | betadisper | 22 | 2 | 1.06 | 0.36 |
| 2015 | adonis | 22 | 2 | 1.05 | 0.31 |
| 2015 | betadisper | 22 | 2 | 0.02 | 0.98 |
| 2016 | adonis | 22 | 2 | 1.63 | 0.14 |
| 2016 | betadisper | 22 | 2 | 4.35 | 0.03 |

**Section A1.3 Figures**

**Figure A1-1.**
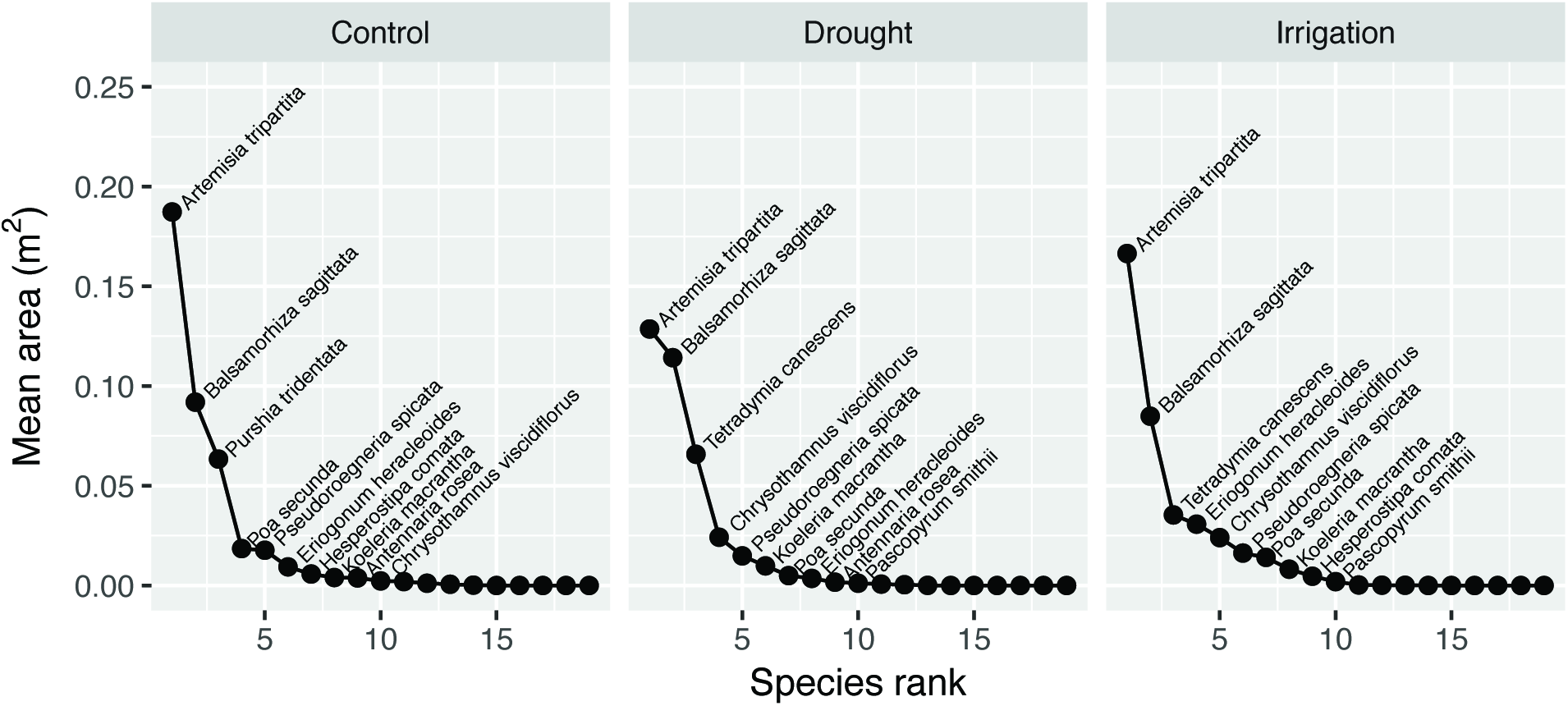
Rank abundance curves for perennial species. Area of individuals (either canopy or basal area cover, depending on life form) was summed withnin years and plots, and then the total area values were averaged across years and plots for each treatment.

**Figure A1-2.**
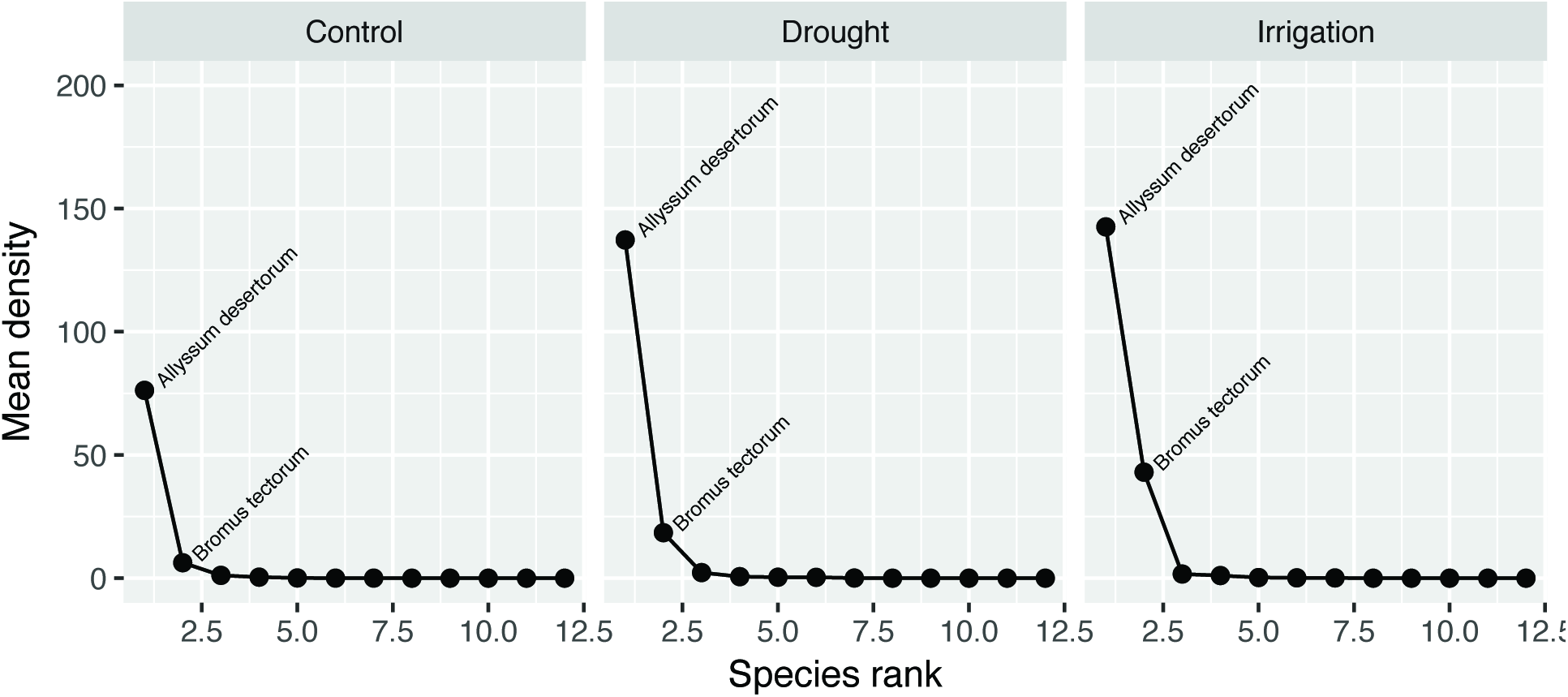
Rank abundance curves for annual species. Density of individuals is averaged across years and plots for each treatment.

**Figure A1-3.**
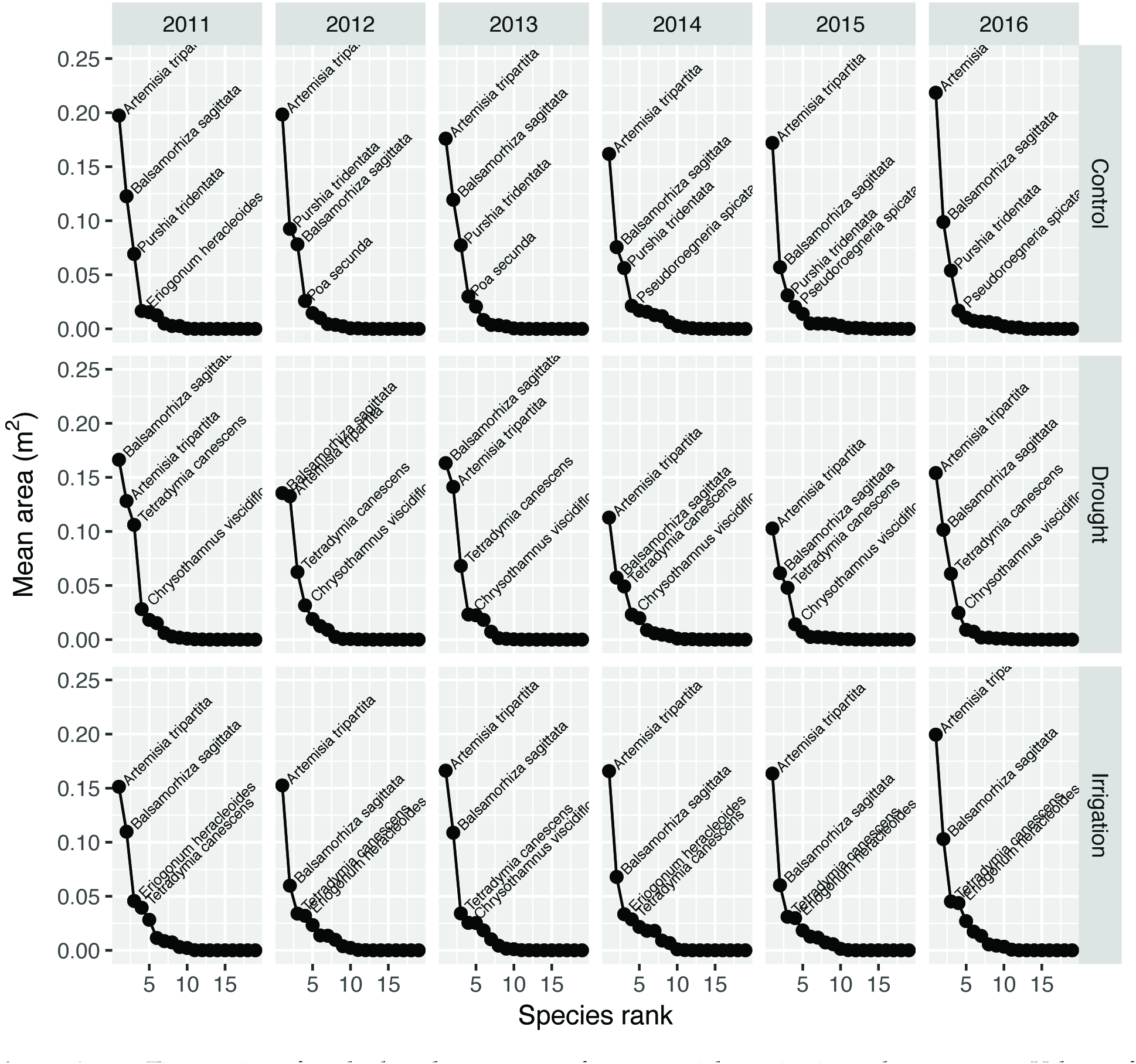
Time series of rank abundance curves for perennial species in each treatment. Values of mean area were averaged over plots. The four most dominant species are labelled in each panel. Area is measured as either basal area or canopy cover area, depending on life form.

**Figure A1-4.**
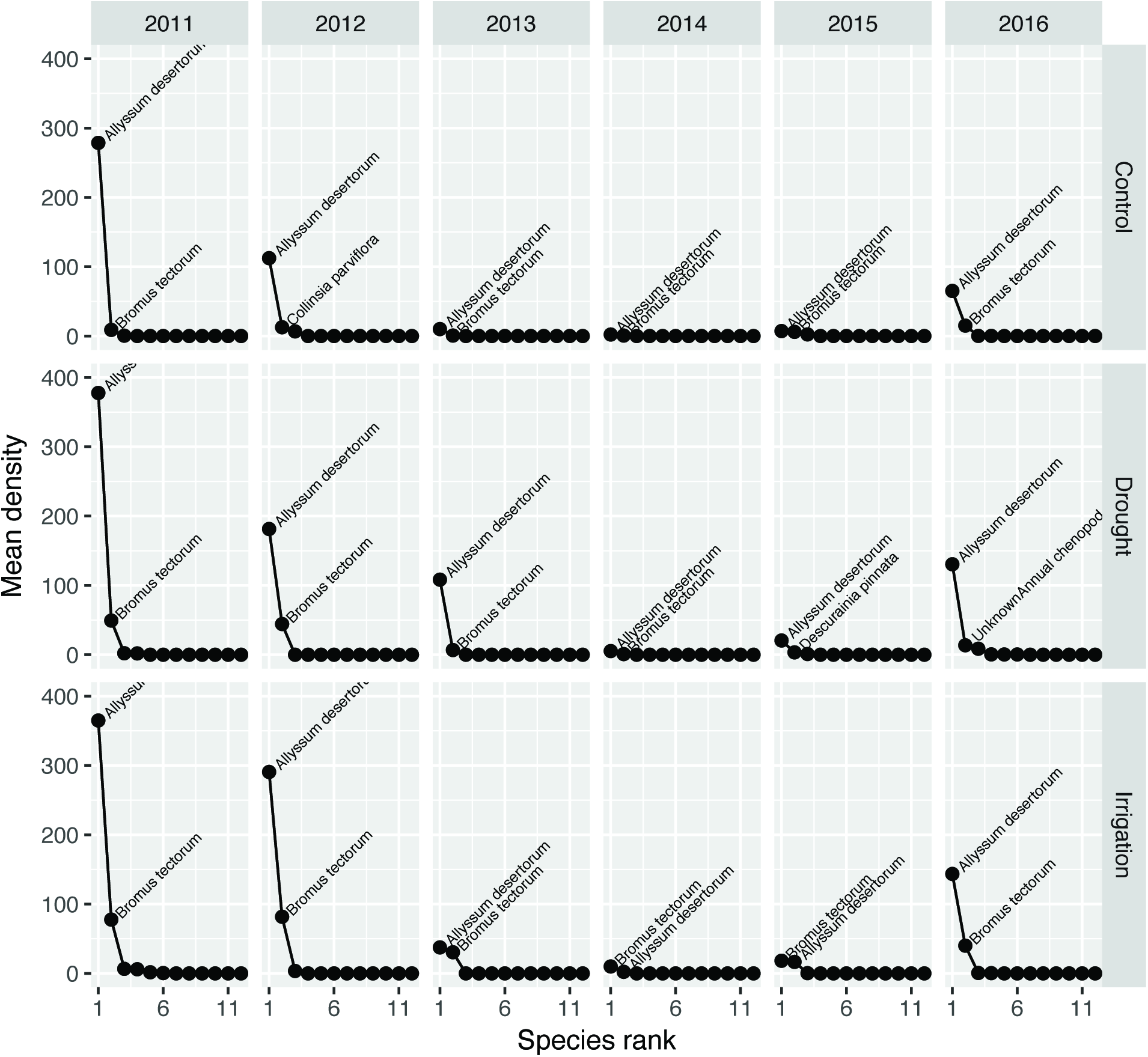
Time series of rank abundance curves for annual species in each treatment. Values of density were averaged over plots. The two most dominant species are labelled in each panel.

**Figure A1-5.**
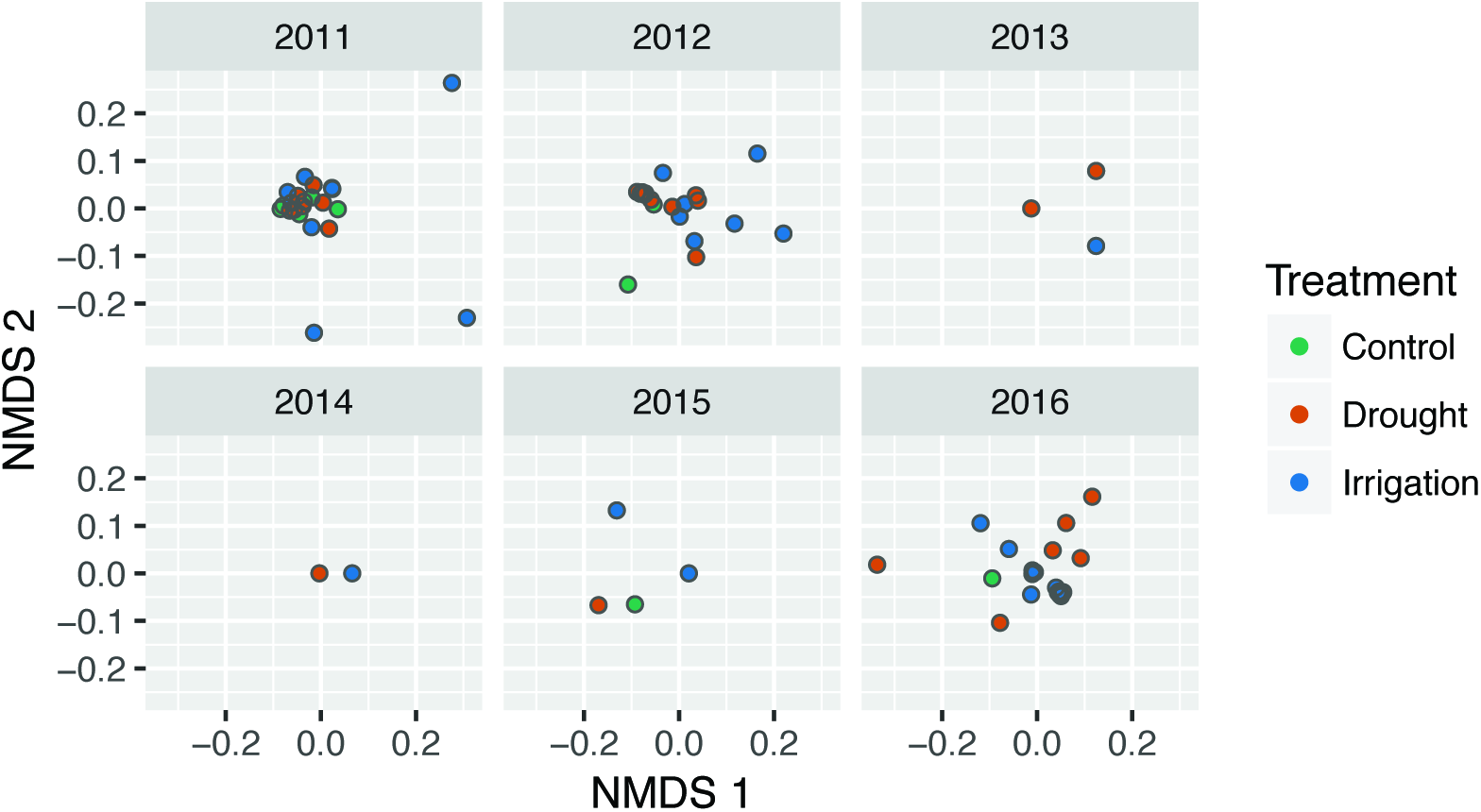
Nonmetric multidimensional scaling scores representing annual plant communities in each plot, colored by treatment.

## Appendix 2 A.T. Tredennick, A.R. Kleinhesselink, J.B. Taylor & P.B. Adler “Consistent ecosystem functional response across precipitation extremes in a sagebrush steppe”. PeerJ

**Section A2.1 Details on SOILWAT predictions**

We used a version of the SOILWAT soil moisture model (Sala et al. 1992) that has been developed specifically for use in semi-arid shrubland ecosystems (Bradford et al. 2014). SOILWAT uses daily weather data, ecosystem specific vegetation data, and site specific soil properties to estimate water balance processes. Specifically, SOILWAT uses daily rainfall data to estimate rainfall interception by plants, evaporation of intercepted water, snow melt and redistribution, infiltration into the soil, percolation through the soil, evaporation from bare soil, transpiration from each soil layer, and drainage. We parameterized SOILWAT using the generic sagebrush steppe parameters and local soil data (Kleinhesselink 2017). SOILWAT was forced by daily weather data collected at the USDA-ARS Sheep Experimental Station over the course of our experiment.

SOILWAT generates soil moisture predictions at several soil depths. We averaged the daily predictions from the upper 40 cm of soil. These predictions represent ambient conditions, similar to our control plots. To generate soil moisture data for our treatment plots, we applied the statistical model described in the main text, which was also used to estimate treatment conditions from control conditions. The time series of those predictions, along with our observations and statistical estimates, is shown in Figure A2-1.

**Figure A2-1.**
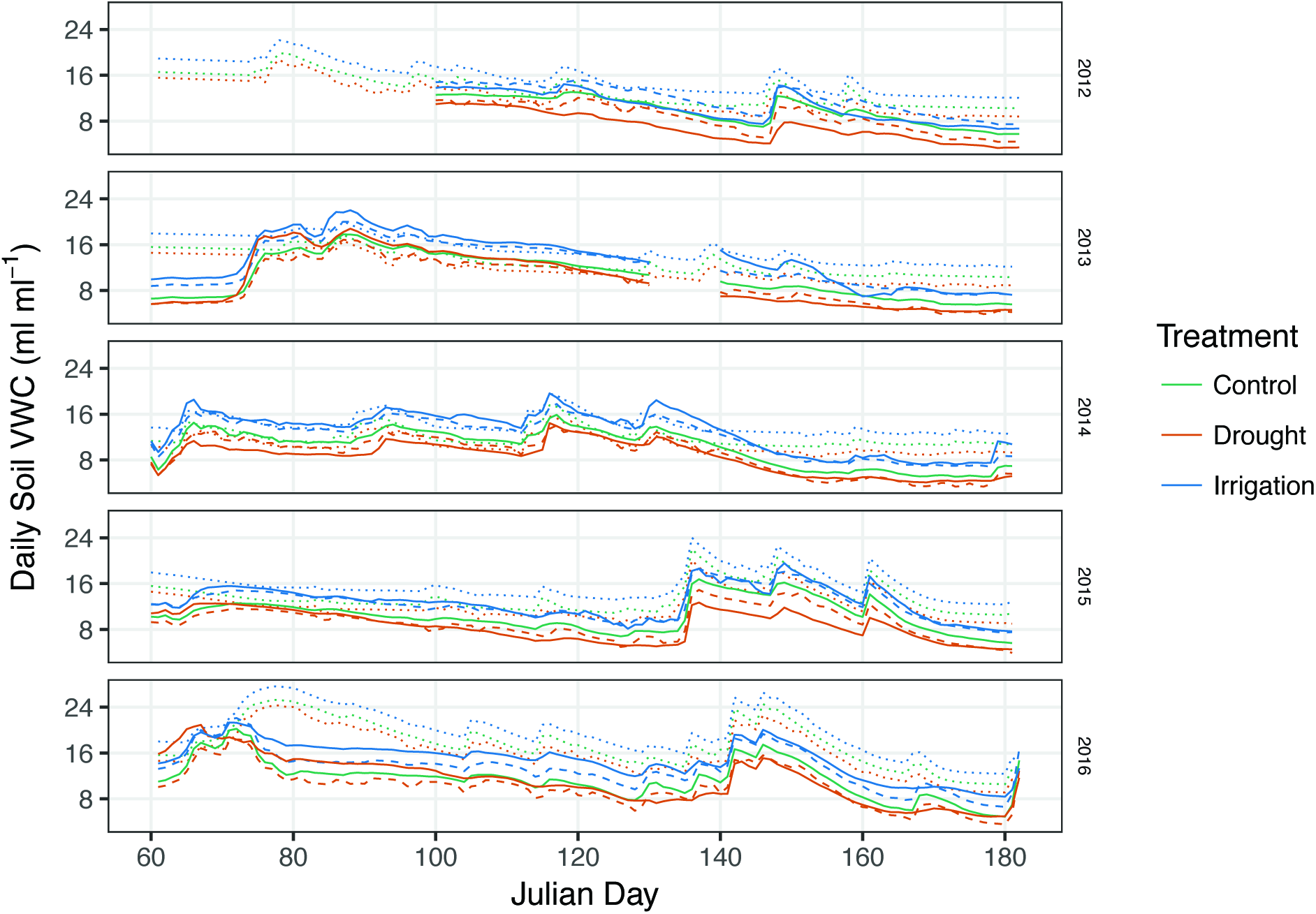
Time series of volumetric water content from March to June in each year from the observed measurements (solid lines), statistical estimates (dashed line), and SOILWAT (dotted line).

## Appendix 3 A.T. Tredennick, A.R. Kleinhesselink, J.B. Taylor & P.B. Adler “Consistent ecosystem functional response across precipitation extremes in a sagebrush steppe”. PeerJ

**Section A3.1 Estimating ANPP**

We used a radiometer to nondestructively estimate aboveground net primary productivity. Our approach relies on relating greenness in a plot to aboveground biomass. In each year we recorded ground reflectances at four bands, two associated with the red spectrum and two associated with the near-infrared spectrum (Table A3-1). We took four readings per plot that were averaged for each band. Bands 1 and 3 correspond to wavelengths collected by the MODIS satellite and bands 2 and 4 correspond to wavelengths collected by the AVHRR satellite.

**Table A3-1.** Radiometer specifications.

| Band number | Spectrum name | Center wavelength | Corresponding satellite |
| --- | --- | --- | --- |
| 1 | red | 626 nm | AVHRR |
| 2 | red | 652 nm | MODIS |
| 3 | near-infrared | 875 nm | AVHRR |
| 4 | near-infrared | 859 nm | MODIS |

Using the RED and NIR reflectance values, we calculate the normalized difference vegetation index (NDVI) for each plot based on both AVHRR-and MODIS-based wavelengths. We calculated NDVI as:

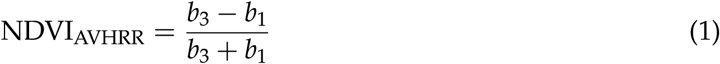

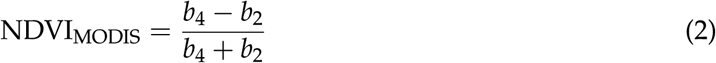

where *b*_*x*_ refers to band *x* (*x* = 1,2,3,4) in Table A1-1.

To convert plot NDVI to biomass, we regressed known biomass values from calibration plots against NDVI calculate for those plots. Calibration plots were located near our experiment plots, and each year we located a new set of 12-16 plots in which we clipped all aboveground biomass, dried it to a constant weight at 60° C, and the weighed. We used these biomass values to estimate regression parameters for both AVHRR-and MODIS-based NDVI. We assessed model fit using *R*^2^ and, for each year, we used the regression parameters associated with the best fit model to estimate biomass in the experimental plots based on their NDVI values (Table A3-2). R code for this procedure is in the file “01_calibrate_radiometer_by_year.R” in the code set.

**Table A3-2.** Details of regression models used to estimate biomass each year.

| Year | Intercept | NDVI Slope | $R^2$ | Min(biomass) | Max(biomass) | $N$ | Algorithm |
| --- | --- | --- | --- | --- | --- | --- | --- |
| 2012 | 9.03 | 144.23 | 0.59 | 8.57 | 41.42 | 15 | MODIS |
| 2013 | 1.44 | 111.39 | 0.39 | 8.63 | 77.62 | 15 | MODIS |
| 2014 | 16.31 | 222.38 | 0.63 | 14.61 | 62.30 | 15 | MODIS |
| 2015 | -84.54 | 386.91 | 0.71 | 44.72 | 129.03 | 12 | AVHRR |
| 2016 | 14.15 | 493.85 | 0.72 | 50.16 | 163.70 | 16 | MODIS |

## Appendix 4 A.T. Tredennick, A.R. Kleinhesselink, J.B. Taylor & P.B. Adler “Consistent ecosystem functional response across precipitation extremes in a sagebrush steppe” PeerJ

**Section A4.1 Random Slopes, Random Intercepts Model**

**Section A4.1.1 Model Description**

Our hierarchical Bayesian model allows us to test for differences among treatments in the relationship between ANPP and soil moisture, and allowed us to account for the non-independence of observations through time within a plot. Treatment differences are modeled as fixed effects, which are modified by plot-level random effects. In what follows, **X** is the fixed effects design matrix including:

1. a column of 1s for intercepts
2. a column of continuous values for the scaled volumetric water content for each observation
3. a column of 0s or 1s indicating whether the observation is from a drought treatment
4. a column of 0s or 1s indicating whether the observation is from an irrigation treatment
5. a column of continuous values for the interaction between scaled volumetric water content and the drought treatment indicator (column 2)
6. a column of continuous values for the interaction between scaled volumetric water content and the irrigation treatment indicator (column 3)

An example of six rows of the fixed effects design matrix (three control plots and three drought plots) is as follows:

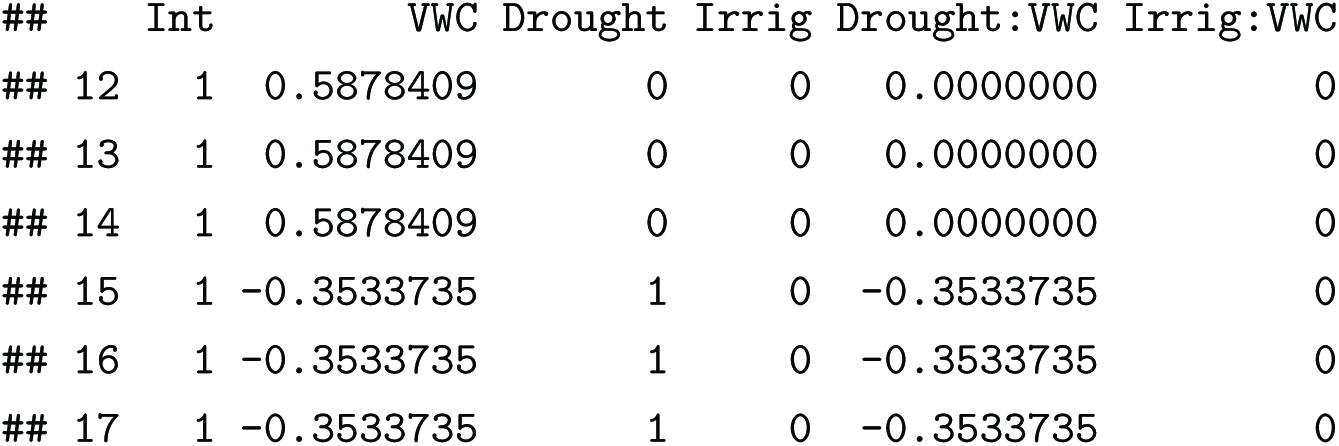

Note that within a year, all plots within a treatment share the same value of volumetric water content. This is because we could not monitor soil moisture in each plot, and instead used sparse observations to model average soil moisture in each treatment in each year (see main text). Using this design matrix, we can estimate six fixed effects (*β*s):

1. the intercept of the soil moisture-ANPP relationship for control plots
2. the slope of the soil moisture-ANPP relationship for control plots
3. the intercept offset for drought plots
4. the intercept offset for irrigation plots
5. the slope offset for drought plots
6. the slope offset for irrigation plots

We are particularly interested in the slope offsets, because these allow us to test whether the slopes for drought or irrigation are different from the control slope. If the slope offset is different from zero, this indicates the slopes are different. We assess whether the slope offsets are different from zero by calculating the probability that the posterior distribution is less or greater than zero (one-tailed tests). If the probability is greater than 0.95, then there is strong evidence that the slope offset is different from zero, and thus different from the control treatment slope.

To account for the fact that observations within plots through time are not independent, we include random effects that modify the fixed effects in each plot. We model these random effects (*γ*s) as offsets drawn from a multivariate normal distribution with mean 0 and a variance-covariance matrix (Σ) that includes a covariance between the intercept and slope offsets. We implement this random effects structure by including a random effects design matrix (**Z**) with a column of 1s for intercept offsets and a column of continuous values for the volumetric water content for each observation.

Lastly, to account for unkown variation across years, we include random year effects. These year effects (*η*s) act as offsets on the intercept.

Putting it all together, our model is defined mathematically as follows, where *i* denotes observation, *j* denotes plot, and *t* denotes year. We assume the observations are conditionally Gaussian,

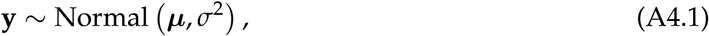

where *μ* is the determinstic expectation from the regression model,

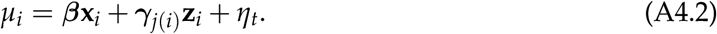

All fixed effect βs were drawn from normally distributed priors with mean 0 and standard deviation 5: *β* ~ Normal(0,5). *γ* random effects were drawn from a multivariate prior centered on zero with a shared variance covariance matrix:

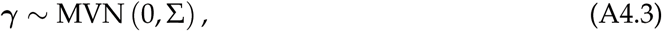

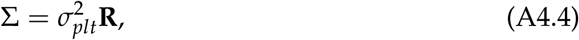

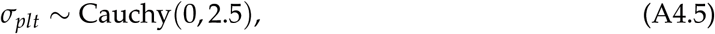

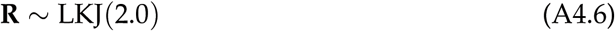

The random year effects (77) are modeled as intercept offsets centered on zero with a shared variance (*σ*_*yr*_): γ ~ Normal (0,*σ*_*yr*_). Σ is the variance-covariance matrix for the intercept and slope offsets for each plot, which is defined as the among plot variance 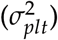 times the matrix **R** that defines the correlation between intercept and slope offsets.

The Bayesian posterior distribution of our model can be expressed as:

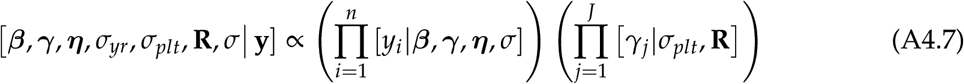

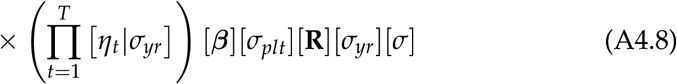

We fit the model using MCMC as implemented in the software Stan (Stan Development Team 2016). Our Stan code is below. All code necessary to reproduce our results has been archived on Figshare (*link here*)and released on GitHub (https://github.com/atredennick/usses_water/releases/v0.1).

**Section A4.1.2 Stan Code**

data {

int<lower=0> Npreds; *# number of covariates, including intercept*

int<lower=0> Npreds2; *# number of random effect covariates*

int<lower=0> Nplots; *# number of plots*

int<lower=0> Ntreats; *# number of treatments*

int<lower=0> Nobs; *# number of observations*

int<lower=0> Nyears; *# number of years*

vector[Nobs] y; *# vector of observations*

row_vector[Npreds] x[Nobs]; *# design matrix for fixed effects*

row_vector[Npreds2] z[Nobs]; *# simple design matrix for random effects*

int plot_id[Nobs]; *# vector of plot ids*

int treat_id[Nobs]; *# vector of treatment ids*

int year_id[Nobs]; *# vector of year ids*

}

parameters {

vector[Npreds] beta; *# overall coefficients*

vector[Nyears] year_off; *# vector of year effects*

cholesky_factor_corr[Npreds2] L_u; *# cholesky factor of plot random effect corr matrix*

vector[Npreds2] beta_plot[Nplots]; *# plot level random effects*

vector<lower=0>[Npreds2] sigma_u; *# plot random effect std. dev.*

real<lower=0> sd_y; *# treatment-level observation std. dev.*

real<lower=0> sigma_year; *# year std. dev. hyperprior*

}

transformed parameters {

vector[Nobs] yhat; *# vector of expected values*

vector[Npreds2] u[Nplots]; *# transformed plot random effects*

matrix[Npreds2,Npreds2] Sigma_u; *# plot ranef cov matrix*

Sigma_u = diag_pre_multiply(sigma_u, L_u); *# plot-level covariance matrix*

for(j in 1:Nplots)

u[j] = Sigma_u * beta_plot[j]; *# plot random intercepts and slopes*

*# regression model for expected values (one for each plot-year*)

for (i in 1:Nobs)

yhat[i] = x[i]*beta + z[i]*u[plot_id[i]] + year_off[year_id[i]];

}

model {

\#### PRIORS

sigma_u ~ cauchy(0,2.5)

sigma_year ~ cauchy(0,2.5)

year_off ~ normal(0,sigma_year); *# priors on year effects, shared variance*

beta ~ normal(0,5); *# priors on treatment coefficients*

L_u ~ lkj_corr_cholesky(2.0); *# prior on the cholesky factor which controls the*

*# correlation between plot level treatment effects*

for(i in 1:Nplots) beta_plot[i] ~ normal(0,1); *# plot-level coefficients for intercept and slope*

\#### LIKELIHOOD

for(i in 1:Nobs) y[i] ~ normal(yhat[i], sd_y); *# observations vary normally around expected values*

}

generated quantities{

corr_matrix[Npreds2] R = multiply_lower_tri_self_transpose(L_u);

cov_matrix[Npreds2] V = quad_form_diag(R,sigma_u);

}

**Figure A4-1.**
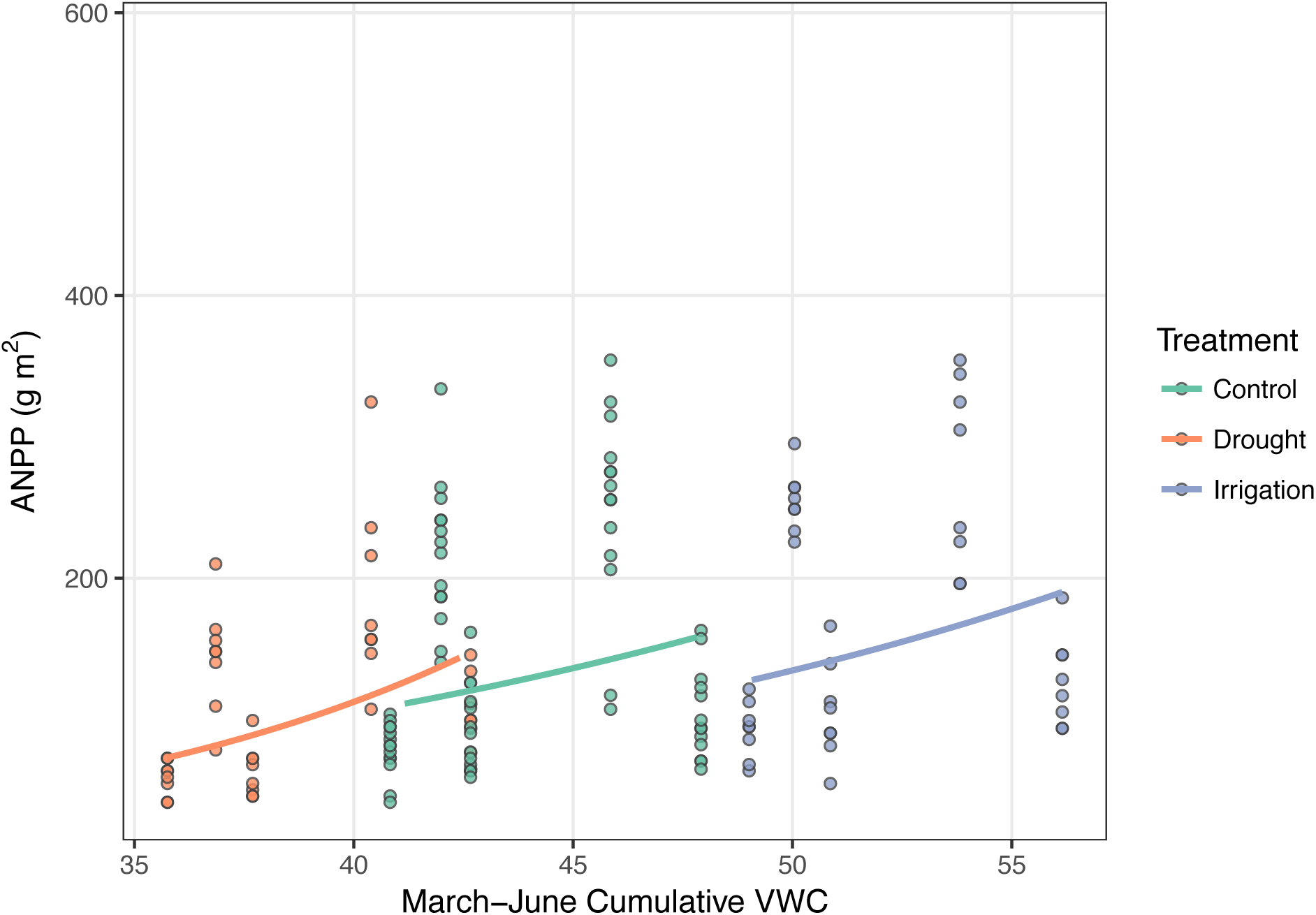
Scatterplot, on the arithmetic scale, of the data and model estimates shown as solid lines. Model estimates come from treatment level coefficients (colored lines).

Stan Development Team. 2016. Stan: A C++ Library for Probability and Sampling, Version 2.14.1.

**Figure A4-2.**
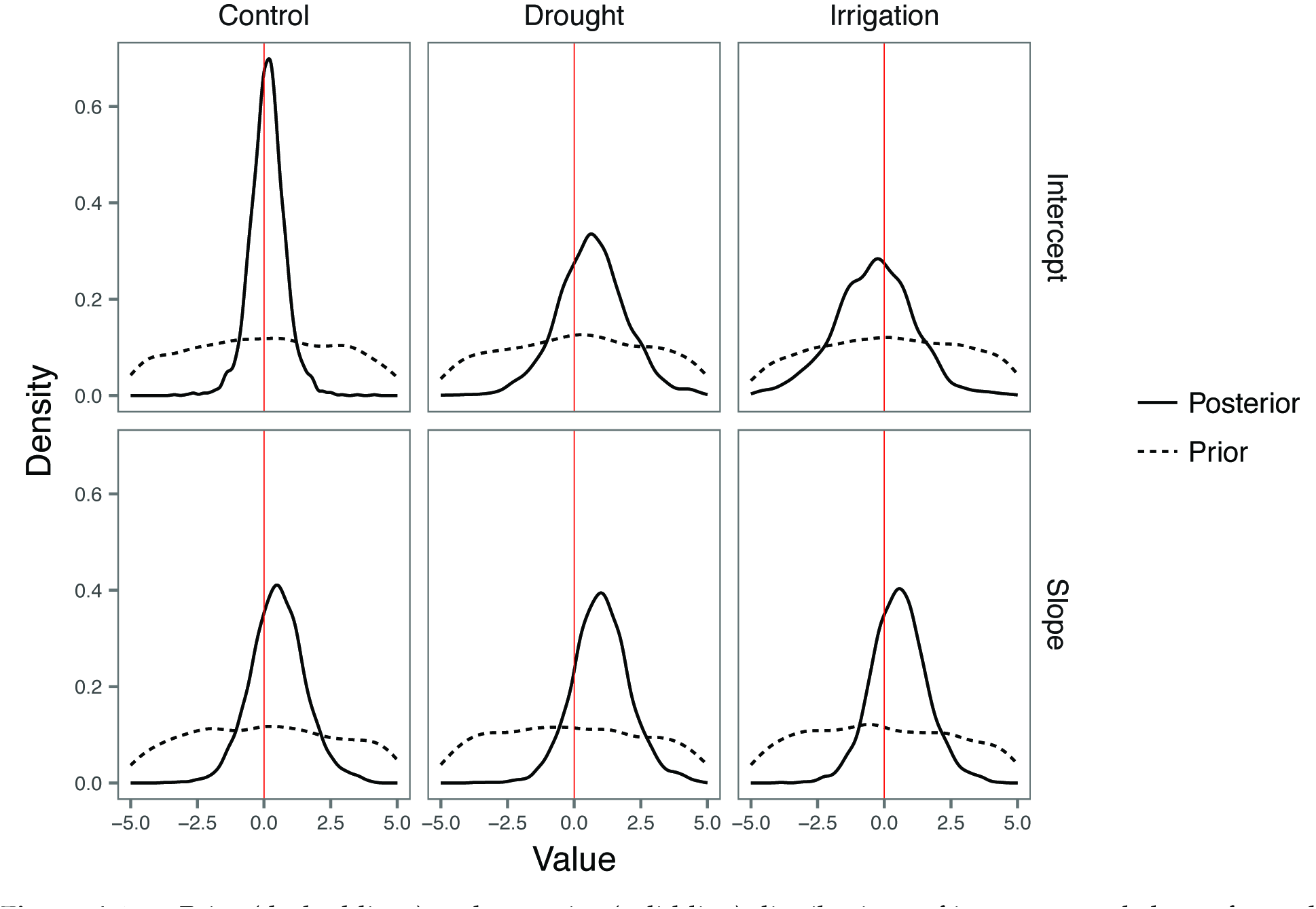
Prior (dashed lines) and posterior (solid line) distributions of intercepts and slopes for each treatment. The slope represents the main effect of soil moisture on log(ANPP). The red line marks 0. Shrinkage of the posterior distribution and/or changes in the mean indicate the data informed model parameters beyond the information contained in the prior for all coefficients.

**Figure A4-3.**
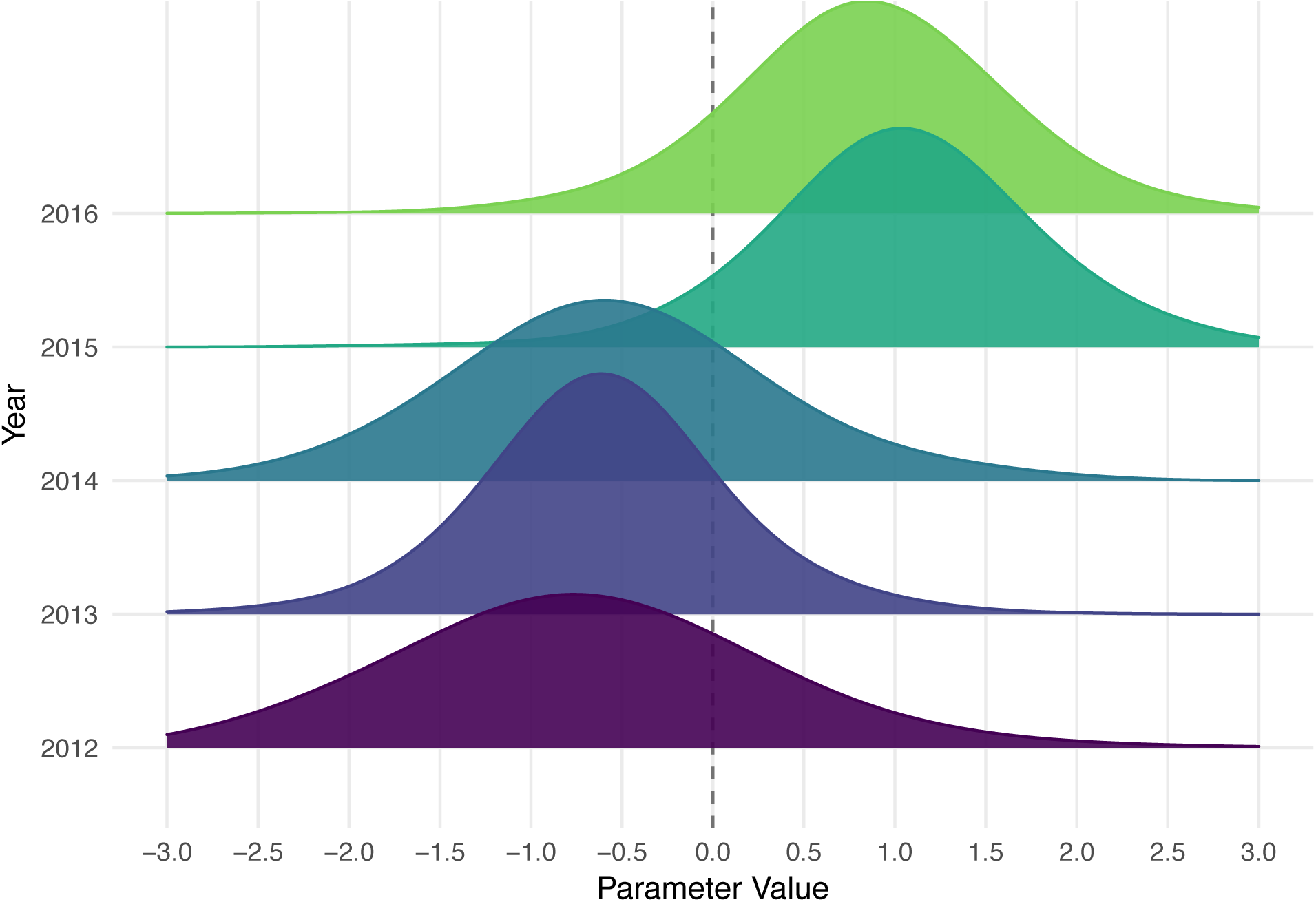
Posterior distributions of random year effects (intercept offsets). Kernel bandwidths of posterior densities were adjusted by a factor of 5 to smooth the density curves for visual clarity.

## Appendix 5 A.T. Tredennick, A.R. Kleinhesselink, B. Taylor & P.B. Adler “Consistent ecosystem functional response across precipitation extremes in a sagebrush steppe” PeerJ

**Section A5.1 Conducting our analysis with NDVI**

The first step of our analysis is to convert NDVI from field-based radiometric measurements to aboveground net primary productivity. However, this conversion is not perfect and the associated uncertainty is sometimes high (Table A1-2). Therefore, we also assessed ecosystem functional response across treatments using NDVI as the response variable rather than ANPP. The model essentially the same as the ANPP model, except we use a Beta likelihood for NDVI since its values range from 0 to 1, not including true 0s or 1s (see Stan code, below).

Our NDVI-model results are very similar to the ANPP-model results (Figure A5-1). As with our ANPP-model results, the positive offset for the drought treatment does not result in a significantly different slope once applied to the control slope (Table A5-1).

**Table A5-1.** Summary statistics from the posterior distributions of coefficients for each treatment (*b* coefficients in equation 1). The ‘Intercept’ and ‘Slope’ summaries reported here for drought and irrigation are from the posterior distributions of the intercept and slope for the control treatment plus the offsets for each treatment. Posterior distributions of the offsets are in Figure A5-1.

| Coefficient | Treatment | Posterior Mean | Posterior Median | Lower 95% BCI | Upper 95% BCI |
| --- | --- | --- | --- | --- | --- |
| Intercept | Control | -1.33 | -1.33 | -2.58 | 0.00 |
| Intercept | Drought | -1.19 | -1.25 | -3.63 | 1.57 |
| Intercept | Irrigation | -1.37 | -1.28 | -4.77 | 1.49 |
| Slope | Control | 0.15 | 0.09 | -1.78 | 2.41 |
| Slope | Drought | 0.48 | 0.43 | -1.53 | 2.87 |
| Slope | Irrigation | 0.18 | 0.13 | -1.70 | 2.50 |

**Figure A5-1.**
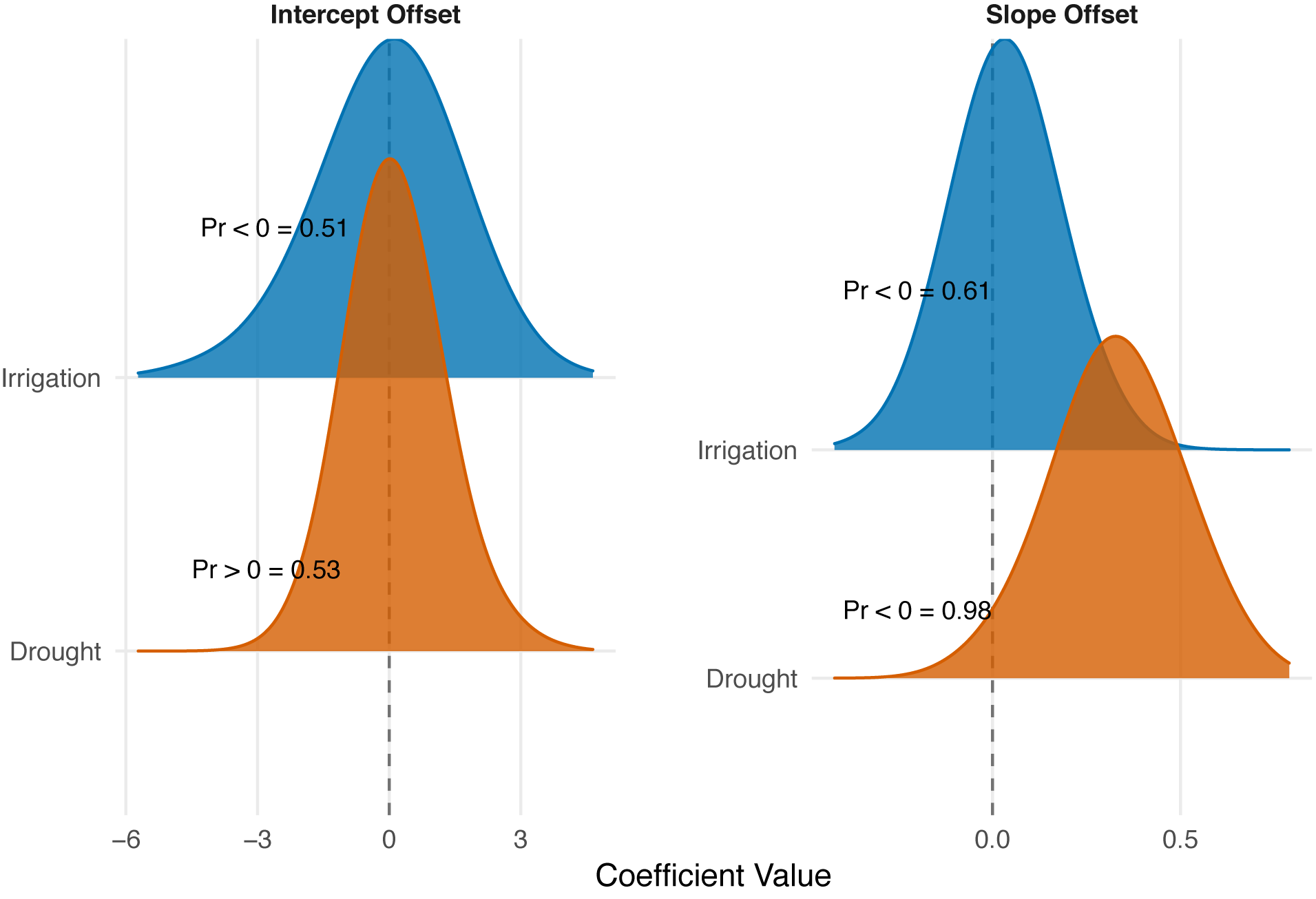
Posterior distributions for the effects of drought and irrigation on the intercept and slope of the soil moisture-NDVI relationship. Treatment effects show the difference between the coefficients estimated in the treated plots and the control plots. Probabilities (Pr </= 0 =) for each coefficient indicate the one-tailed probability that the coefficient is less than or greater than zero, depending on whether the median of the distribution is less than or greater than zero. The posterior densities were smoothed for visual clarity by increasing kernel bandwidth by a factor of five.

**Section A5.1.1 Stan Code**

data {

int<lower=0> Npreds; *# number of covariates, including intercept*

int<lower=0> Npreds2; *# number of random effect covariates*

int<lower=0> Nplots; *# number of plots*

int<lower=0> Ntreats; *# number of treatments*

int<lower=0> Nobs; *# number of observations*

int<lower=0> Nyears; *# number of years*

vector[Nobs] y; *# vector of observations*

row_vector[Npreds] x[Nobs]; *# design matrix for fixed effects*

row_vector[Npreds2] z[Nobs]; *# simple design matrix for random effects*

int plot_id[Nobs]; *# vector of plot ids*

int treat_id[Nobs]; *# vector of treatment ids*

int year_id[Nobs]; *# vector of year ids*

}

parameters {

vector[Npreds] beta; *# overall coefficients*

vector[Nyears] year_off; *# vector of year effects*

cholesky_factor_corr[Npreds2] L_u; *# cholesky factor of plot random effect corr matrix*

vector[Npreds2] beta_plot[Nplots]; *# plot level random effects*

vector<lower=0>[Npreds2] sigma_u; *# plot random effect std. dev.*

real<lower=0> sd_y; *# treatment-level observation std. dev.*

real<lower=0> sigma_year; *# year std. dev. hyperprior*

real<lower=0> phi; *# dispersion parameter*

}

transformed parameters {

vector[Nobs] A; *# parameter for beta distn*

vector[Nobs] B; *# parameter for beta distn*

vector[Nobs] yhat; *# vector of expected values*

vector[Npreds2] u[Nplots]; *# transformed plot random effects*

matrix[Npreds2,Npreds2] Sigma_u; *# plot ranef cov matrix*

Sigma_u = diag_pre_multiply(sigma_u, L_u); *# cholesky factor for plot-level covariance matrix*

for(j in 1:Nplots)

u[j] = Sigma_u * beta_plot[j]; *# plot random intercepts and slopes*

*# regression model for expected values (one for each plot-year*)

for (i in 1:Nobs) yhat[i] = inv_logit(x[i]*beta + z[i]*u[plot_id[i]] + year_off[year_id[i]]);

A = yhat * phi;

B = (1.0 - yhat) * phi;

}

model {

\#### PRIORS

phi ~ cauchy(0, 5);

sigma_year ~ cauchy(0,2.5);

sd_y ~ cauchy(0,2.5);

year_off ~ normal(0,sigma_year); *# priors on year effects, shared variance*

beta ~ normal(0,5); *# priors on treatment coefficients*

L_u ~ lkj_corr_cholesky(2.0); *# prior on the cholesky factor which controls the*

*# correlation between plot level treatment effects*

sigma_u ~ cauchy(0,2.5);

for(i in 1:Nplots) beta_plot[i] ~ normal(0,1); *# plot-level coefficients for intercept and slope*

\#### LIKELIHOOD

y ~ beta(A, B); *# observations vary according to beta distribution*

}

generated quantities{

corr_matrix[Npreds2] R = multiply_lower_tri_self_transpose(L_u);

cov_matrix[Npreds2] V = quad_form_diag(R,sigma_u);

}

## Appendix 6 A.T. Tredennick, A.R. Kleinhesselink, J.B. Taylor & P.B. Adler “Consistent ecosystem functional response across precipitation extremes in a sagebrush steppe” PeerJ

**Section A6.1 Characterizing Extreme Precipitation Amounts**

Following the proposed methods of Lemoine et al. (2016), we calculated quantiles from the empirical distribution of growing season precipitation at Dubios, ID. We chose the 1% quantile to be indicative of extreme dry conditions (drought) and the 99% quantile to be indicative of extreme wet conditions (irrigation). The data consist of 91 yearly records, which we assume are approximately normally distributed for these purposes. The R code below shows our procedure, and Fig. A6-1 shows the results.

\## Water year defined as precip in Oct-Dec in year *t* and Jan-Sept in year t+1

\## following USGS.

first_water_months <- c(′10′,′11′,′12′) *# first months in water year, to be promoted a year*

weather <- read.csv(“‥/data/weather/dubois_station_weather_01092018.csv”) %<%

dplyr∷select(DATE, PRCP) %<%

dplyr∷rename(“date” = DATE, “precip” = PRCP) %<%

separate(date, into = c(“year”, “month”, “day”), sep = “-”) %<%

mutate(precip = ifelse(is.na(precip), 0, precip)) %<% *# set missing station data to 0*

mutate(year = as.numeric(year)) %<%

mutate(water_year = ifelse(month %in% first_water_months, year+1, year)) %<% *# create water years*

filter(year != 1925) %<% *# remove first year because don’t have first water-year months*

group_by (water_year)%<%

summarise(annual_precip = sum(precip)) %<%

rename(year = water_year)

mean_ppt <- mean(weather$annual_precip)

quants_ppt <- quantile(weather$annual_precip,probs = c(0.01,0.99))

quants_ppt[1]/mean_ppt*100 *# percent of mean ppt for drought*

\## 1%

\## 26.38659

quants_ppt[2]/mean_ppt*100 *# percent of mean ppt for irrigation*

\## 99%

\## 183.5835

ggplot(weather, aes(x=annual_precip))+

geom_histogram(bins=20, color=“dodgerblue”, fill=“dodgerblue”, aes(y=‥density‥))+

geom_line(stat="density”, color="blue”)+

geom_vline(aes(xintercept=quants_ppt[1]), linetype=2)+

geom_vline(aes(xintercept=quants_ppt[2]), linetype=2)+

ylab(“Density”)+

xlab(“Growing Season Precipitation (mm)”)+

theme_bw()+

theme(panel.grid.minor = element_blank())

**Figure A6-1.**
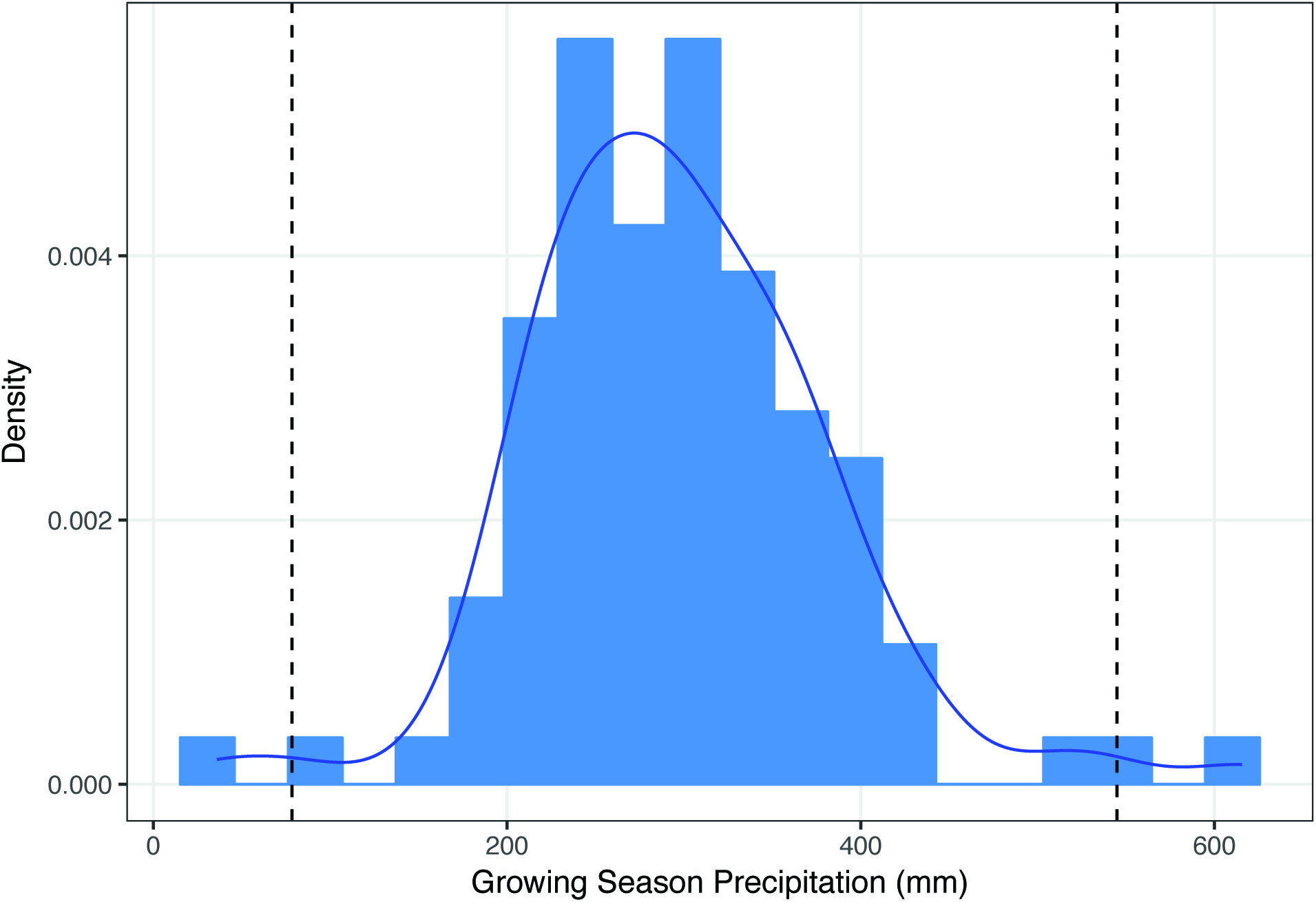
Density of the empirical distribution of growing season precipitation at Dubois, ID. Dashed vertical lines show the 1% and 99% quantiles, assuming a normal distribution.

## Footnotes

1 Dates formatted as: mm-dd-yyyy.

